# Environmental perturbation increases gene expression variability and unmasks genetic regulation for transcriptional robustness

**DOI:** 10.64898/2026.02.18.706644

**Authors:** James Phipps-Tan, Saudat Alishayeva, Huiting Xu, Julien F. Ayroles, Luisa F. Pallares

## Abstract

Environmental perturbation reshapes phenotypes not only by shifting trait means but also by modulating phenotypic variability. Whether environmentally induced changes in variability -environmental robustness- are genetically regulated remains unclear. To address this question, we collected nearly 2,000 matched genomes and transcriptomes from an outbred population of *Drosophila melanogaster* before and after exposure to a high-sugar diet, and quantified how dietary stress remodels expression variability and its genetic architecture. We find that environmental perturbation induces a transcriptome-wide increase in expression variability, with the notable exception of key developmental pathways. At the same time, stress unmasks pervasive genetic regulation for expression variability (veQTL). These loci are mostly environment-dependent, show distinct functional and evolutionary constraints from mean-regulating loci, and appeared detrimental when they confer to much robustness to dietary perturbation. Together, our findings establish gene expression variability as a central dimension of the organismal response to environmental changes, and uncover its genetic basis as a distinct, evolutionary constrained layer of phenotypic regulation beyond mean effects.

## Main text

The extent to which environmental conditions shape phenotypic variation and its underlying genetic architecture remains a central problem in evolutionary biology. Much progress has been made in understanding plastic responses (i.e., changes in mean trait value)^1^ and their genetic basis ^2–4^. However, although research on robustness and canalization has shown that phenotypic variability (i.e., the dispersion of individuals around the trait mean) also responds to environmental perturbation^5–7^, the degree to which environmental robustness is under genetic control remains mostly unexplored.

In contrast with plasticity-regulating loci, robustness-regulating alleles differ not in their mean phenotype but in their degree of phenotypic variability. Biological noise theory has shown that variability-regulating loci might be beneficial when populations are far from the fitness peak, e.g., when exposed to new environments^8,9^. And, if the degree of variability correlates with fitness, these alleles can themselves be targets of selection^8,10,11^. However, empirical progress has been limited because identifying environment-dependent loci that regulate phenotypic variability requires large individual-level datasets across multiple controlled environmental conditions.

Here, we tackle this question by focusing on gene expression variability, a trait that has been extensively studied in several species^12–16^. Although some variability-regulating loci have been identified ^17–20^, the global architecture of the regulation of expression variability, including its pervasiveness across the transcriptome and its sensitivity to environmental perturbation, remains mostly uncharacterized.

To address this, we collected nearly 2,000 individual genomes and transcriptomes from an outbred *Drosophila melanogaster* population and identified loci that regulate genotype- and environment-dependent expression variability (Fig. 1A). Flies were exposed to two environmental conditions, the ancestral laboratory diet and a novel high-sugar diet which negatively affects metabolism, development, and lifespan^21,22^. We show that dietary stress induces a transcriptome-wide increase in expression variability, while only a subset of development-related pathways remain highly robust. This change in variability was associated with abundant cryptic genetic variation, where most variability-regulating loci (veQTL) were detected in the high sugar diet and showed minimal signal under the standard diet. Notably, veQTLs showed stronger signatures of negative selection compared to mean-regulating eQTLs. In particular, high-robustness alleles are kept at low frequencies in this population, suggesting that low expression variability may be disadvantageous in novel environments.

**Figure 1:**
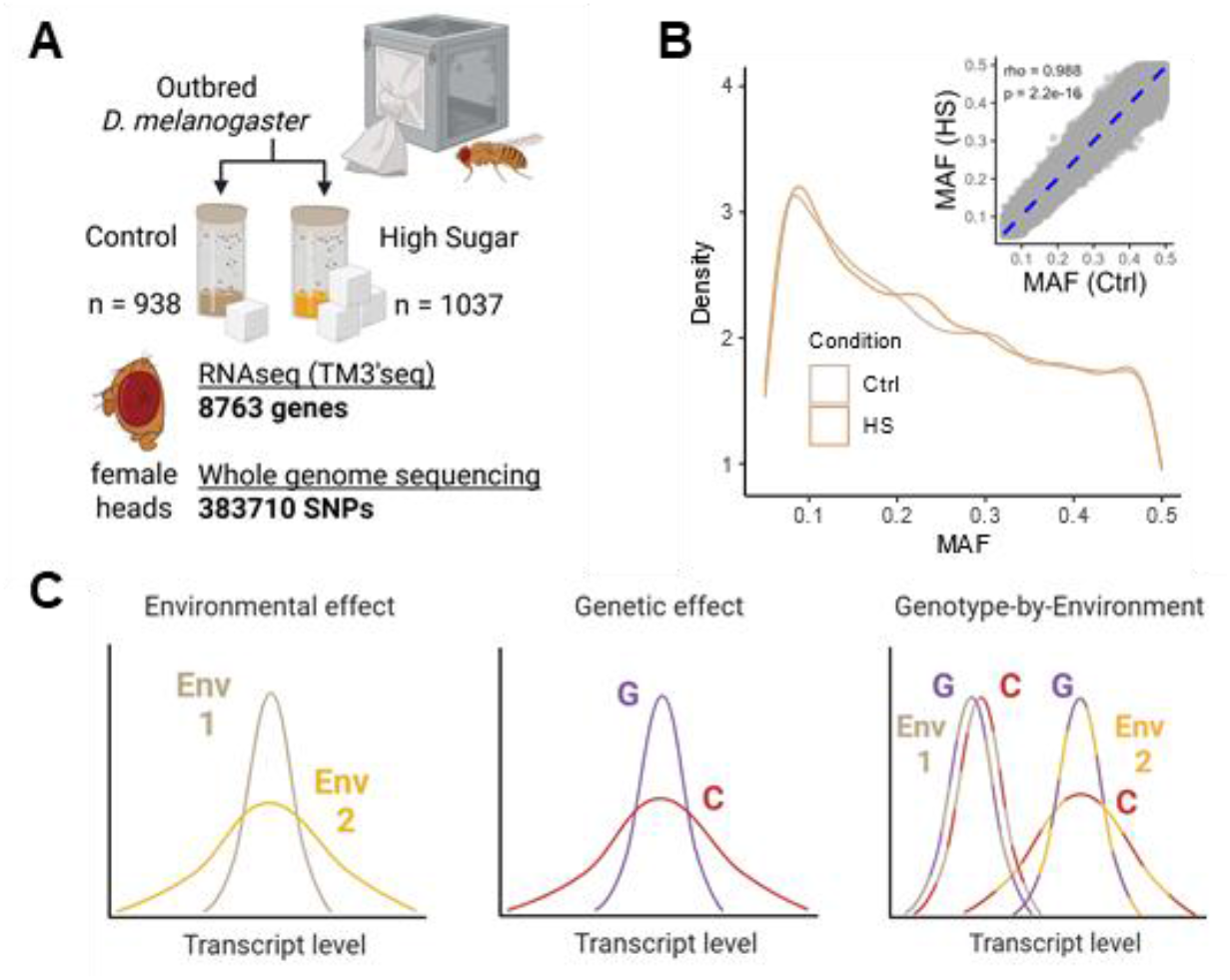
Experimental design to explore the environmental and genetic regulation of transcript level variability. **(A)** A *Drosophila melanogaster* outbred synthetic population (Dros-OSP) from the Netherlands was exposed to two dietary conditions (control: 8% sugar, high sugar: 20% sugar). Eggs developed into adults in each condition, and 7-day old females were used for RNA and DNA extraction. RNAseq and whole-genome DNA libraries for each one of the 1975 flies were prepared following the TM3’seq protocol^25^. The final dataset consisted of 8763 expressed genes and 383 710 SNPs (MAF > 0.05, LD < 0.8). **(B)** The site frequency spectrum (density kernel = 2) and the minor allele frequencies (MAF) are highly correlated between both diets (p = 2.2 e -16, rho = 0.99). The dashed blue line represents the best fit linear regression. **(C)** Depiction of an environmental effect (left), genetic effect (middle) and genotype x environment (GxE) effect (right) on transcript level variability where the variability-increasing effect of allele C is only manifested under environment 2. Differences in mean transcript level were depicted to better visualize the GxE effect, but GxE effects might affect expression variability without affecting the mean. (A) and (C) were created in BioRender. Phipps-Tan, J. (2025) https://BioRender.com/0gs1ie0 and https://BioRender.com/i4wuzfv

Together, these results show that environmental perturbation not only modifies the degree of phenotypic variability in a population, but also reshapes the genetic architecture for gene expression, exposing to selection cryptic variability-regulating loci when environmental shifts occur.

### Dissecting the effect of Genotype, Environment, and GxE interactions on expression variability

To assess the transcriptional response to environmental perturbation, we collected eggs from one of the *Drosophila Outbred Synthetic Populations* (Dros-OSP) derived from the Netherlands^23^, and let them develop for one entire lifecycle, from egg to adult, either on standard (8% sugar) or high sugar media (20% sugar). We collected 7-day-old females and generated head transcriptomes and genomes for 1,975 flies, 938 and 1,037 from standard and high sugar diets, respectively (Fig. 1A, fig. S1, S2). Using 383,710 SNPs genotyped in at least half of the 1,975 flies (MAF > 0.05, LD R^2^ < 0.8, fig. S2), we show that the allele frequency spectrum is highly correlated between diets (rho = 0.99, Fig. 1B), and that there is no diet-related population structure in a genomic PCA (fig. S2D). This indicates that gene expression differences between diets are not due to genetic differentiation, allowing us to dissect how environmental perturbation, genotype, and genotype-by-environment interactions shape expression variability (Fig. 1C). We then quantified the effect of high sugar on gene expression variability at two levels. At the transcriptome-wide level, we asked whether the variability hierarchy of the most and least variable genes was altered in high sugar. At the level of individual genes, we asked whether the amount of expression variability differed in the two conditions for specific genes.

### Diet-independent functional and evolutionary constraints shape variability ranks within the transcriptome

Gene-expression variability differs widely among genes, and across species the most and least variable genes are often conserved^12–16,24^. Whether environmental perturbation alters this transcriptome-wide organization remains poorly characterized. To test this, we quantified gene-wise expression variability using the median absolute deviation (MAD) computed from variance-stabilized expression values, and compared ranks between standard and high sugar conditions (Fig. S5, table S1, S2). Variability ranks were strongly correlated between diets (Spearman correlation, rho = 0.895, p-value < 2.2e-16, Fig. 2A), suggesting that environment-dependent factors only minimally disturb the variability structure of the transcriptome, and that gene-specific functional and evolutionary properties are the main determinant of differences between genes.

**Figure 2:**
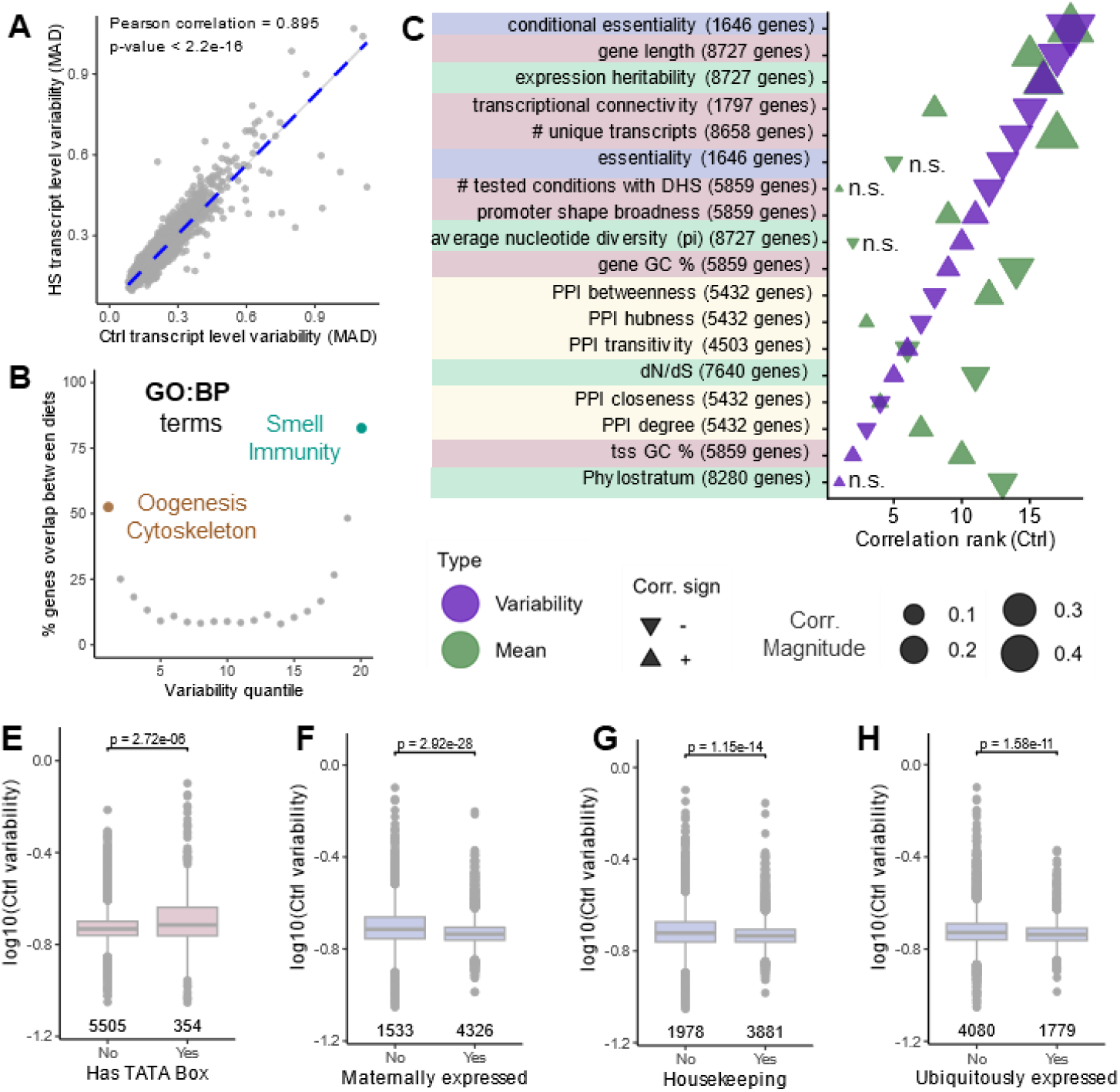
Gene-wise transcript level variability correlates with gene regulatory features, gene function and sequence evolution. (A) Gene-wise transcript level variability under the high-sugar diet is highly correlated with variability under the control diet. Each dot is one gene out of the 8763 genes in the transcriptome. (B) Out of all 5% quantiles in each diet, the lowest- (brown) and highest-variability (cyan) genes overlap the most, at 52% and 77% respectively. The significant Gene Ontology:Biological Process (GO:BP) terms estimated for the overlapping genes are highlighted for low and high variability quantiles (g:Profiler adjusted p-value <0.05). (C) Gene-level molecular and evolutionary features correlate with the level of expression variability, and are different for average expression values. Here we show the results for control diet, but the results are the same for high sugar data (fig. S6). Features were clustered by their type: red = transcriptional control, yellow = protein function, blue = higher-order function, green = sequence diversity and evolution. Significance of the correlation was assessed by partial Spearman correlations while adjusting for mean expression levels, and using Spearman correlations for mean expression. All features except those indicated with n.s. are significant (table S3). (E-H) Discrete features that correlate with transcript level variability. (E) TATA box genes have higher variability, while (F) maternally-expressed, (G) housekeeping and (H) ubiquitous-expressed genes have lower variability (difference in medians and p-values of Wilcoxon rank-sum tests shown). Number of genes indicated above ‘Yes/No’ labels. For C-H, variance-stabilized (VST) count matrices were used to compute MAD and averages for each gene. See table S5, and fig. S6 and S7 for additional correlates.

The overlap across diets was strongest at the extremes of the expression variability hierarchy. 77% of the top most variable, and 50% of the least variable genes overlap between diets (top = 5^th^, bottom = 95^th^ variability percentiles, Fig. 2B). This exceeds the 5% expected overlap of random gene sets of equivalent size drawn from two ranked lists based on a hypergeometric distribution. The most variable genes in the *Drosophila* transcriptome are enriched for ‘response to external stimuli’ and ‘immunity’ GO terms, as reported in mice, humans, and plants^12,14,16^ (Hypergeometric test, adjusted p-value < 0.05, Fig. 2B, table S3). On the other hand, the least variable genes have very specific enrichments in ‘reproduction’, ‘gamete development’, and ‘microtubule cytoskeleton organization’ (Hypergeometric test, adjusted p-value < 0.05, Fig. 2B, table S4). This contrasts with previous studies using single cell data or biopsies of non-reproductive tissue^14–16^ that have linked low-variability genes to cellular housekeeping functions. Here, we quantified expression variability in a body part -the fly head-that regulates oogenesis^25–27^ which allowed us to uncover essential organismal-level functions that are strongly constrained to maintain low expression variability.

We then asked which factors explain the expression variability rankings. For this, we compiled gene-level annotations spanning regulatory architecture, gene structure, network connectivity, essentiality, and evolutionary constraint, and estimated their correlation with gene-level expression variability (table S5). Across both diets, high-variability genes tend to be at the periphery of protein-protein interaction networks, and are involved in fewer protein-protein interactions (Fig. 2C, fig. S6), consistent with stronger constraint on variability for genes embedded in multi-protein complexes, or densely connected modules^14,24^. Higher variability was also correlated with longer genes and more spliced variants (Fig. 2C, fig. S6), suggesting that transcription or translation duration could impact transcript level variability^28^. More variable genes also tend to have promoter and chromatin features previously linked to greater transcriptional stochasticity, including broader promoters, higher GC content around transcription start sites and across gene bodies, increased DNase hypersensitivity, and the presence of a TATA box in the promotor (Fig. 2D–H; Figs. S6– S7). Conversely, hallmarks of functional constraint like housekeeping roles, essentiality or conditional essentiality, ubiquitous expression, and maternal transmission, were features correlated with low-variability genes (Fig. 2, E-H, fig. S7).

Because we used an outbred fly population where expression differences among individuals reflect both micro-environmental^29^ and genetic effects^30^, we also asked whether genetic variation features were correlated with expression variability ranks. As reported for mammalian systems^12,14^, high-variability genes in *Drosophila* tend to have greater nucleotide diversity (pi), higher dN/dS values, and higher heritability (Fig 2C; fig. S6). In this species, these genes also tend to have more eQTLs and veQTLs (fig. S8, S9 and table S5).

### Dietary stress induces transcriptome-wide increase in gene expression variability

We then asked whether high sugar affects the degree of expression variability of individual genes. For this, we jointly tested for diet effects on mean expression and expression variability using GAMLSS, which models differential mean and differential variance in a unified framework^28^. We find a transcriptome-wide variability response, with 88% of the genes showing significant differences between high sugar and standard diets, while only 50% showed differential mean expression (FDR 5%, Fig. 3A, table S7 and S8, fig. S10-12).

**Figure 3:**
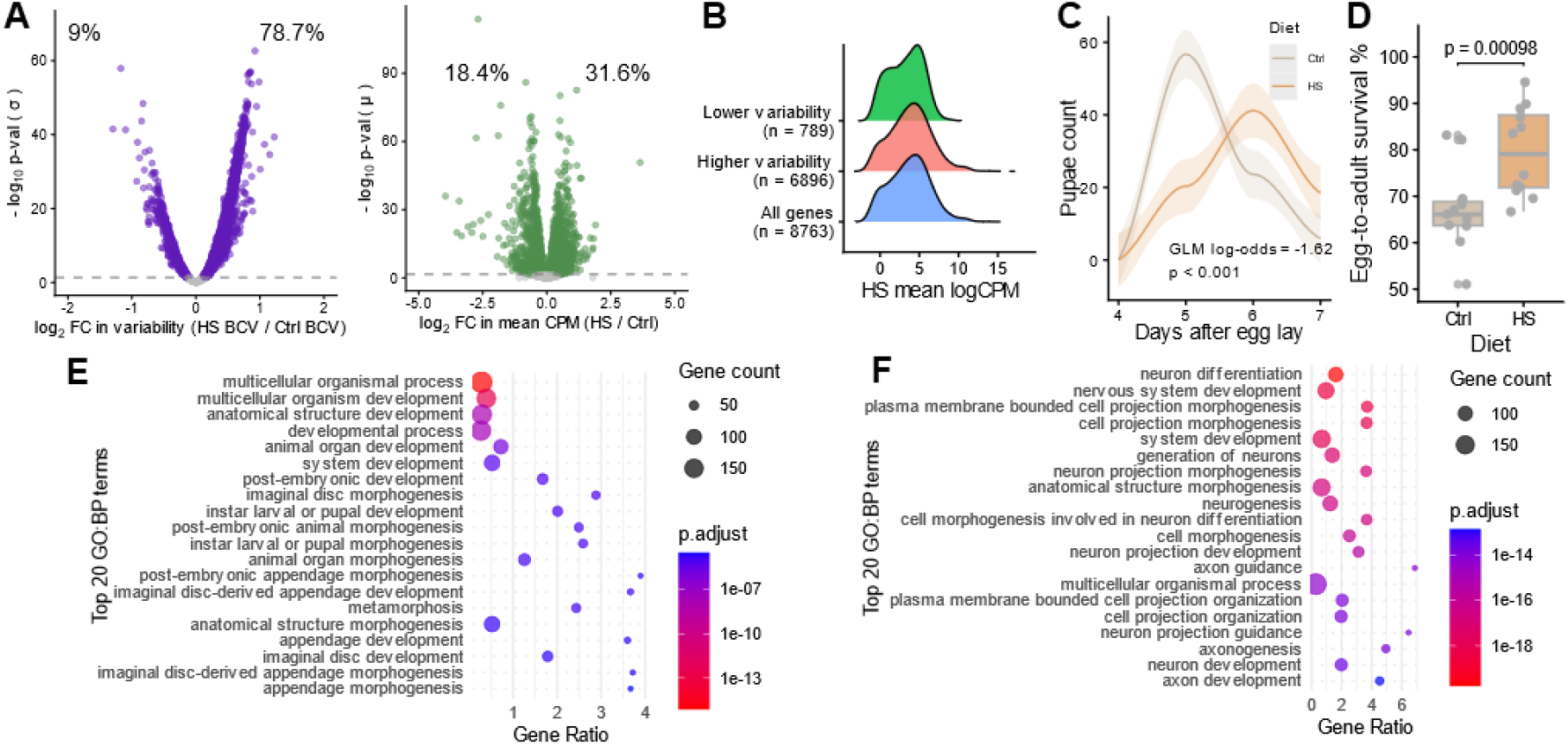
High sugar modifies transcript level variability in a large fraction of the transcriptome. (A) Volcano plot showing in purple the genes with significant changes in transcript level variability (biological coefficient of variation, BCV. GAMLSS FDR 5%). And in green, the genes with significant changes in average gene expression (mean CPM. GAMLSS FDR 5%). The dotted grey line depicts FDR 5% threshold. The percentages indicate the fraction of the transcriptome that increase/decrease mean/variability. (B) The average expression level for the whole transcriptome is shown, as well as for the genes that increase or decrease expression variability in high sugar conditions. The genes that are detected as differentially-variable between conditions span the entire range of average expression levels of the population. (C-D) Diet-dependent development time and survival in the Dros-OSP Netherlands outbred population. Compared to equivalent control flies, flies from the high-sugar population (C) take longer to pupate (discrete binomial generalized linear model log-odds = -1.62, p < 0.001) and (D) have higher egg-to-adult survival (Wilcoxon rank-sum test p = 0.00098, median difference = 13%) (E) Top 20 GO biological process enrichments for genes that decreased in variability but did not change in mean. (F) Top 20 GO biological process enrichments for genes that decreased in variability and changed in mean. Gene ratio is calculated as (intersection size/GO term size)/(GO term size/8763). p.adjust is calculated by a hypergeometric test corrected for multiple testing using the default g:SCS procedure.

Remarkably, most of the differentially-variable genes became more variable under the novel high sugar condition (90%, Fig. 3A). This increase was independent of mean expression, spanned all expression levels, and occurred regardless of the direction of mean change (Fig. 3B and fig. S11). This pattern suggests either a systemic dysregulation of transcription in response to environmental perturbation^4–6^, or the existence of a central on-off switch for transcriptional noise reminiscent of the role of heat-shock proteins in exposing phenotypic noise^29,31^. Genes that increase in variability without a detectable change in mean (n = 3444, 39% of the transcriptome) have a very specific enrichment for ‘RNA processing’ and ‘RNA metabolism’ GO terms (hypergeometric test, adjusted p-value<0.05, table S9), suggesting RNA regulation as a candidate mechanism linking dietary stress to increased expression variability.

Contrary to the transcriptome-wide trend, a small subset of genes became less variable in response to high sugar. These genes are enriched for “development”, “metamorphosis” (390 genes with only variability changes; 4.5% of the transcriptome, hypergeometric test, adjusted p-value < 0.05, Fig. 3E, table S10), and “neurogenesis” GO terms (399 genes with both, variability and mean changes; 4.6% of the transcriptome, Fig. 3F, table S11).

While relating gene expression variation to organismal phenotypic variation remains a major challenge, these results motivated us to examine whether metamorphosis itself shows any indication of tighter regulation under high sugar. For this, we measured developmental time and egg-to-adult viability (i.e., number of eclosing adults/number of eggs). We find that in high sugar, eggs take significantly longer to develop as reported before^21^ (20% delay, discrete time GLM, p-value = 0.001, Fig. 3C; table S12), but strikingly, they develop into adult flies with a higher viability rate than in standard conditions (13% increase, Wilcoxon rank-sum test p-value = 9.8e-4, Fig. 3D; fig. S13; table S12). How tighter regulation of these developmental genes contributes to increased egg-to-adult viability requires further experimental exploration. However, these results suggest an active sugar-sensitive mechanism linking reduced expression variability to lengthened, but not compromised, development.

### Robustness of gene expression to dietary stress is under genetic control

To determine if robustness to environmental perturbation has a genetic basis, we asked whether genotypes differ in the degree of expression variability when exposed to high sugar (Fig. 4A). For this, we mapped variance-eQTLs (veQTL) using a squared residual correlation test implemented in veqtl-mapper^32^ after mapping and accounting for genetic effects on mean expression (eQTL) identified with tensorQTL^33^. Modelling explicit genotype-by-environment interactions for genome-wide veQTL mapping is not currently computationally practical. Thus, we mapped QTLs separately for each diet, calling a QTL significant at FDR 5% (table S13).

**Figure 4:**
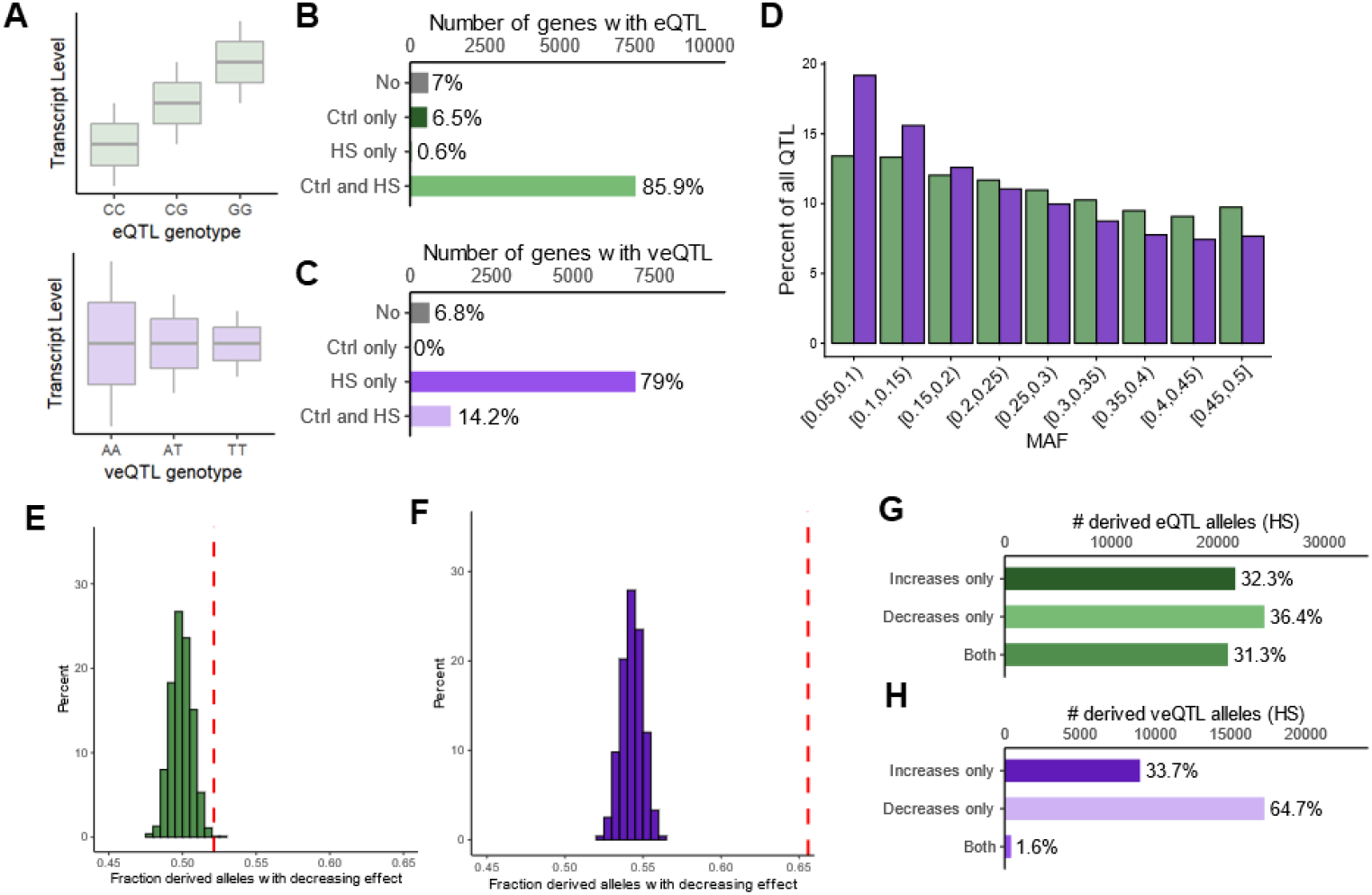
Variability-regulating genetic effects are environment-dependent and variability-reducing alleles tend to be detrimental. (A) Cartoon depiction of genotypes conferring different transcript level means (eQTL) or variabilities (veQTL). (B-C) Number of and proportion of all genes with no, GxE (control-only, HS-only), and QTL in both diets. Results are shown for (B) eQTL and (C) veQTL. GxE determined by FDR 5% in one condition and 10% in the other. (D) The minor allele frequency (MAF) distribution of veQTL is biased toward rare alleles (purple; n=111116, median = 0.211) compared to eQTL (green; n=306334, median = 0.249) suggesting stronger negative selection (Kolmogorov Smirnoff test p<2.2e-16, Wilcoxon ranked-sum text p< 2.2e-16). (E – H) Different properties of derived alleles at trans-eQTL and trans-veQTL in high sugar. The results for control diet follow the same patterns (fig. S22). (E-F) Fraction of QTL derived alleles that decrease expression variation (red dotted line = median fraction derived from 1000 subsamples) compared to the null distribution derived from 1000 random subsamples of 5000 MAF-matched non-significant SNPs (green and purple histograms) (table S18). (E) 51% of eQTL derived alleles decrease mean expression level, representing the 0.002 percentile of the null distribution. (F) 67% of veQTL derived alleles decrease expression variability. (G) Effect of the eQTL derived allele on expression levels. An allele can always increase, always decrease, or both decrease and increase expression levels depending on the focal gene. When pleiotropic, eQTL alleles can have different effects on the target genes, suggesting mean-regulation effects are gene specific. (H) Effect of the veQTL derived allele on expression levels. An allele can always increase, always decrease, or both decrease and increase transcript level variability depending on the focal gene. When pleiotropic, veQTL alleles have the same effect on all focal genes, suggesting that variability-regulation effects are SNP-specific.

To quantify the extent of genotype-by-environment interactions, we defined a QTL as having a shared effect across diets if it was detected at 5% FDR in one diet and at 10% FDR in the other. This definition is conservative for shared effects because small-effect QTLs might fall below the detection threshold in only one diet, yet it provides a principled summary of environment-dependence at the transcriptome scale. We find that while most genes have both, eQTLs (96%, n = 8431) and veQTLs (93%, n = 8167), there are different environmental-specificities. Genes tend to have eQTL regulation across diets (86% genes; Fig. 4B), while veQTL regulation is detected predominantly in high sugar (14% both diets, 79% only high sugar, Fig. 4C). The low proportion of shared veQTL effects is robust to other significance thresholds, heterozygote frequency, and data normalization (fig. S14-S15, table S13-S15). These results show that expression variability is under widespread environment-dependent genetic control, with most veQTL effects being unmasked only under high sugar.

The genetic architecture for expression variability is dominated by trans-effects (veQTL more than 10kb away from the focal gene), with only 1,6% of the genes having detectable cis-veQTLs (n =136). This pattern was also observed in mice, where environment-dependent veQTLs were predominantly trans-acting^34^. The degree to which veQTL effects are driven by direct single-locus effects or through higher order interactions (e.g., epistatic or genotype-by-microenvironment) requires further investigation^17,32^. Nonetheless, the dominant trans-architecture of variability could be brought about by a larger mutational target, trans-acting modulators of variability and network-level dysregulation.

Each veQTL was associated with a median of 1-2 genes, suggesting low pleiotropy (table S13), yet, a small fraction was associated with more than 40 genes (2% of veQTL, table S16, fig. S16 and S17). The more pleiotropic of these putative expression variability hotspots seem to regulate genes involved in RNA processing and RNA metabolism (fig. S18; table S16), pointing to a potential role of the RNA processing machinery in regulating overall expression variability.

### Low-variability alleles appear disadvantageous in novel environments

Given such widespread genetic control for gene expression variability, the question arises of whether variability-regulating loci are subject to selection. Theoretical work predicts that high-noise alleles (i.e., high-variability alleles) can be beneficial when populations are far away from the fitness peak^8,9^. However, high phenotypic variability has also been associated with detrimental effects^15,35^. Empirical progress has been limited due to the lack of genome-wide veQTL catalogs. Here, we approach this question in a comparative framework where variability-regulating loci (veQTL) are compared to mean-regulating loci (eQTL) because there is no a priori expectation that high-expression alleles could be generally detrimental or beneficial, while there are clear predictions of the effects of high-variability alleles. We focus on trans-acting QTLs because there are very few cis-veQTL (but see results for all QTL types in table S17).

First, we compared allele frequency and functional properties of QTLs. We find that, in stark contrast with eQTLs, veQTLs are significantly depleted in promoters and exons, in particular in nonsense and missense mutational effects, while overrepresented in introns and intergenic regions (table S17, chi-squared tests). Accordingly, veQTLs also show a stronger bias toward low allele frequencies than eQTL and background SNPs (median MAF: veQTL = 0.21, eQTL = 0.25, all SNPs = 0.23, Kolmogorov Smirnoff test p<2.2e-16; Wilcoxon rank-sum test p< 2.2e-16; Fig 4D; fig. S19). These results, together with recent findings in human cellular traits^36^ and yeast expression^37^, support an evolutionary model where genetic variation that modulates variability is subject to stronger purifying selection than mean-regulating loci.

Next, we tested whether high-variability alleles show signatures that could indicate a general detrimental or beneficial effect. We reason that, based on the nearly-neutral theory of evolution^38^ that states that recent genetic variation tends to be slightly deleterious, any bias in variability effects of derived alleles could be indirectly associated with detrimental effects in this population. Towards this, we determined the ancestral state of 121,683 SNPs in the experimental *D. melanogaster* population by comparing its genetic polymorphisms to that of two sister species, *D. yakuba* and *D. simulans*.

Our results show that derived veQTL alleles tend to decrease expression variability, and this bias is more pronounced when their effect is assessed in high sugar (66% bias in HS, 57% bias in Ctrl; Fig. 4E; fig. S20B; table S18). As expected, derived eQTLs and background alleles show a very small bias, if any (52%, Fig. 4E, F; fig. S20A; table S18). Consistent with stronger constraints on low-variability alleles, they are maintained at lower frequency than high-variability derived alleles, particularly in high sugar (median DAF low-variability = 0.20, median DAF high-variability = 0.69, Wilcoxon rank-sum test p-value < 2.2e-16, fig. S21, table S19). These results are robust to sample size and MAF differences with background SNPs (table S20). In addition to this bias in allelic effects, veQTL alleles show directional effects across genes, that is, alleles are either high- or low-variability alleles (Fig. 4G; fig. S20D; table S18). On the other hand, eQTL alleles can be high- or low-expression depending on the focal gene (Fig. 4H; fig. S20C; table S18).

Together, our results challenge the assumption that high-variability alleles are generally detrimental, and suggest that for gene expression, alleles that are highly robust to environmental perturbation (i.e., low-variability) might be especially disadvantageous when populations are exposed to new environmental conditions^8–10^.

## Discussion

Our finding that a novel stressful environment results in an increase in phenotypic variability and the unmasking of cryptic genetic effects is consistent with robustness and canalization theory^5,6^. Yet, from the first experiments done by Waddington^39,40^, the new phenotypes observed in and selected for under environmental perturbation were associated with mean-regulating cryptic loci. Here we show that the exploration of new phenotypic space in stressful conditions goes beyond mean control, and that variability-regulating loci also play a critical role.

Gene-specific functional and evolutionary factors constrain the amount of expression variability that a gene exhibits ^13–16^. Here we show that within such gene-level constraints, in an outbred population, expression variability between individuals is genetically regulated. Although the sources of non-genetic expression variability have been extensively discussed and quantified, in particular in unicellular organisms ^29,30^, the mechanisms behind genetically-encoded expression variability remain mostly unknown. Two of our results point towards RNA metabolism and processing as potential mediators of transcriptome-wide expression variability. Further experiments based on reverse genetics are needed to validate whether mutations in genes that regulate these processes do indeed generate changes in transcriptome-wide variability, while single-cell transcriptomics promise to resolve such molecular processes to cellular resolution.

Extraordinary progress has been made in understanding the genetic basis of gene expression variation by focusing on allelic effects on mean expression (eQTL). But here we have shown that expression variability, an overlooked aspect of expression variation, is also genetically regulated and its genetic architecture shows distinct evolutionary constraints than mean-regulating loci. These results imply that to gain a full understanding of the genetic basis of complex traits -and therefore of human disease, phenotypic prediction, and forecasting of adaptation to climate change-phenotypic variability needs to be considered a bonafide trait instead of just a nuisance in phenotypic distributions.

## Acknowledgements

We specially thank members of the Ayroles lab for help during fly collection and processing of the genomics data. Thanks to P. Andolfatto for help with the ancestral state inference. We thank A. Gecele, V. Sacks-Rattenbach and D. Korkmaz for helping with the experiments to estimate developmental time and egg-to-adult survival. We thank J. S. Lotharukpong and J. Barrera-Redondo for support and advice on gene age and OrthologR, and N. Kalábová for recommending programmatic gProfiler. We thank A. A. Brown for help with using veqtl-mapper. We thank Detlef Weigel and Jun Ishigohoka for critical feedback and helpful suggestions on this manuscript, and all members of the Pallares lab for helpful discussions.

## Author contributions

Conceptualization: J.P-T, J.F.A, L.F.P. Methodology: J.P-T, S.A, J.F.A, L.F.P. Investigation and Bioinformatics Analysis: J.P-T, S.A, H.X. Writing—original draft: J.P-T and L.F.P. Writing— review and editing: J.P-T, S.A, J.F.A, L.F.P. Visualization: J.P-T. Supervision: L.F.P. Funding acquisition: J.F.A, L.F.P.

## Code availability

All code needed to reproduce the analyses, figures, and tables is available at https://github.com/PallaresLab/EnvGen_ExpVar/.

