## Supplementary methods and figures for "Environmental perturbation increases gene expression variability and unmasks genetic regulation for transcriptional robustness"

**The PDF file includes:**

Materials and Methods

Figs. S1 to S21

References

### Materials and Methods

#### Outbred *Drosophila melanogaster* population

We used the Netherlands population (Nex) from the Drosophila Outbred Synthetic Populations, Dros-OP[1] . This population was created by crossing 19 Netherlands-derived inbred lines[2] in a round-robin design before outbreeding for ~125 generations by the time of the experiment, with a population size ~50 000-100 000 individuals. This results in a mapping population that has been used to lab conditions for many generations and where linkage has been broken down by recombination resulting in high statistical power and high resolution for genetic mapping.

#### Experimental design

To assess the effect that environmental perturbation has on gene expression, and the degree to which such effects are driven by genotype-by-environment interactions, we exposed the Nex outbred population described above to two dietary conditions - the control diet used to maintain the population for the last hundred generations, composed of 1% agar, 8.3% glucose, 8.3% yeast, 0.41% phosphoric acid (7%), and 0.41% propionic acid (83.6%), and a high sugar diet with 20% more sugar - 1% agar, 8.3% glucose, 13.3% sucrose, 8.3% yeast, 0.41% phosphoric acid (7%), and 0.41% propionic acid (83.6%). All flies were maintained in the same incubators at 25 °C, 65% humidity, and a 12h:12h light:dark cycle.

Eggs were collected from the outbred population in several alternating rounds of either high sugar or control bottles. In this way, we prevented a scenario where flies would have the choice of laying eggs either on control or on high sugar media, as this could result in genetic structure in the next generation. Eggs developed into adults within each diet and mated seven days old female flies were collected during several days. The eclosion date as well as the round of egg laying were recorded and explored as covariates.

### Fly head sample collection

Fly head sample collection was carried out using the same procedure and apparatus as described in the 'Experimental Design' section in Pallares *et al.*, *Cell Genomics* 2025. In short, female flies were knocked out with ice and placed in 96-well plates in a cold room. The plate was submerged in liquid nitrogen to flash-freeze it, and the plate was banged against a lab bench to separate the head from the body. Fly heads were separated into a new 96-well plate using costume-made sieves while preserving the body-head ID for each individual fly. All flies used are 7-days-old from the date of eclosion. The control flies used in this study were reported in Pallares *et al.*, *Cell Genomics* 2025. The high-sugar flies were collected in the same experiment and at the same time as the control samples, and they are used for the first time here.

### RNAseq library preparation, data processing, filtering and estimation of batch effects

Fly heads in the 96-well plates were processed for mRNA extraction and library preparation following a protocol previously optimized for *Drosophila* single heads (TM3'seq[3])). Raw reads for control and high sugar flies were mapped to the r6.14 *D. melanogaster* genome using Star aligner [4], and feature counts[5] was used to assign reads to the 17727 annotated genes.

Non-expressed and lowly expressed genes were removed for each diet separately, first, by keeping genes with at least 1 CPM in at least 80% of the samples, and an average CPM>1. For this we used TMM library size normalisation[6] and edgeR cpm()[7]. Next, we removed 7 genes mapped to the Y chromosome that presumably mis-mapped due to high sequence similarity with non-Y genes based on FlyBase orthology comparisons. This resulted in 8779 genes expressed in 938 control samples and 1037 high sugar samples.

Due to the scale of the experiment, the data was collected in multiple batches, with control and high sugar samples distributed as evenly as experimentally possible across the batches. To account for their effect on mean transcript level, we first transformed the raw gene count matrix containing both, control and high sugar data, in two ways that best fit the downstream analyses (Fig. S3). First, we used the *voom*[8] function from the package *limma*[9] with normalization factors that account for library size estimated in edgeR [7]. The resulting counts are log(CPM) and normally distributed, showing a negative mean-variability relationship (Fig. S3B). This is used for the QTL mapping and details of the analyses, including how mean effects are accounted for in the veQTL mapping, are described under the 'eQTL and veQTL mapping' section. Second, we used *vst*, the variance-stabilizing transformation from DESeq[10] with the following parameters (blind = F, method = 'parametric', design = ~diet). This results in expression normalized by library size and composition, and are approximately normally distributed where the transcript level variability per gene is more or less independent of mean expression (Fig. S3C). This is used for the correlations between genewise variability and functional and evolutionary features (see details below).

For each matrix, we ran Type-II MANOVA of the first 100 PCs against known experimental batches and diet, followed by pairwise Chi-squared tests and Cramer's correlation, identifying five non-redundant batches with variable but significant effects on mean transcript level - well, plate, library preparation batch, sequencing batch, egg lay batch and plating batch. The significant batch effects were removed from each gene expression matrix using removeBatchEffect(design = ~diet) from the package limma [9]. Visual inspection of PCA plots for each matrix before and after batch effects were removed confirmed that these batch effects indeed accounted for some PC1 structure. However, even when all batch effects were accounted for, there remained residual structure in PC1 not explained by diet. Thus, we identified surrogate variables (SVs) using the function sva()[11] from the package sva to account for this residual structure. Based on PCA inspections following different numbers of surrogate variable removal, we selected 3 and 4 SVs for the VST- and voom-transformed matrices, respectively. These SVs, in addition to the significant experimental batches, were, depending on the analysis, either regressed out of the matrices using removeBatchEffect() (correlation analyses and QTL mapping), or explicitly modelled as batch variables (differential expression mean and variability analysis). Finally, to identify genes with non-unimodal expression distributions, undesired as their intrinsic high-variability will bias our estimation of transcript level variability and affect all subsequent analyses, we applied a Hartigan's dip test to each matrix with batch effects and SVs regressed out and identified 16 genes (two unique to the voom matrix, four unique to the VST matrix and 10 shared) with non-unimodal distributions at a Benjamini-Hochberg procedure adjusted p-value (FDR) of 0.05. We removed these 16 non-unimodal genes, yielding a final set of 8763 genes in both populations. RNAseq quality control metrics are summarised in Fig. S1.

#### Correlation between transcript level variability and molecular features

To evaluate whether certain molecular, functional, and evolutionary gene features were associated with expression variability, we first estimated expression variability for each gene within each dietary condition. For this, we used the variance-stabilized gene count matrix (VST) because this transformation explicitly controls for the mean-variability relationship in RNAseq data. We regressed out the five experimental batches and three surrogate variables using removeBatchEffect() to obtain a batch-free VST matrix. We then split this batch-free VST matrix by diet, and for each gene calculated the median absolute deviation (MAD). MAD per gene is estimated as the median of the absolute distances of each individual expression level from the median expression of all individuals and corresponds to the variability metric per gene.

While greatly reduced relative to the MAD of raw and log(CPM) counts (fig. S3A and S3B), there still exists a mean-variability correlation in the MAD of VST counts (fig. S3C). To ascertain if the imperfect mean-variability correlation was artefactually introduced by the VST transformation, two alternative metrics of variability were calculated (table S1). The coefficient of variation was not used because the mean logCPM of some genes after batch effect correction is negative (Fig. S3B). Adjusted variance, inspired by F* in [12] and adjusted SD in [13], is obtained by dividing the diet-population-specific variances by the average variance-mean relation derived from a degree 5 polynomial regression of the whole-dataset log(variance) against mean. This degree was the highest degree polynomial for which a higher degree polynomial did not significantly explain more variance in the data as determined through the R package anova(). Expression variation, from [14], is obtained by running a local regression of log(variance) against mean for each condition and obtaining the residuals. Following the original procedures, both metrics were estimated using the TMM library size-normalised logCPM matrix with five batches and four SVs regressed out. The persistence of the residual correlations across all three metrics (Fig. S4), alongside a previous report of a similar pattern in *D. melanogaster* embryos [13] suggests that the correlation is not artefactual and property of the *D. melanogaster* transcriptome. This highlights the presence of higher-order mean-variability correlations in transcriptomes which cannot be fully eliminated through simple regression procedures and stresses the need for, where available, further mean-controlling procedures (e.g., partial correlation with mean expression) when trying to identify features predictive of transcriptome-wide variability.

Additionally, outliers and sample size might strongly influence variability estimation. To determine which metric was more robust to outlier effects, we carried out leave-one-out analysis by re-calculating the MAD on the VST matrix with each individual excluded once and assessing the maximum rank change in the variability estimate across each leave-one-out estimate. We calculated the variance, SD, and IQR as well for comparison. Given that the MAD yielded the lowest rank change out of the metrics, it is the least sensitive to single outliers (table S2).

To determine the effect of sample size in variability estimations, we randomly downsampled 10, 50, 100, 200, 300, 500 and 900 individuals from the VST matrix with replacement and calculated the MAD for each gene. We downsampled 100 times per sample size and obtained the average MAD across the 100 downsamples. Then, we calculated the percent difference from the average downsampled MAD values from the MAD estimated on the full sample size. We found that using less than 100 individuals resulted in a genome-wide underestimation of expression variability, and that MADs for many genes deviated well over 10% from the full-sample MAD (Fig. S5). We suggest that a minimum of 100 samples should be used to get a proper estimate of genome-wide variability and accurate estimates of gene-specific variability. In this study, we are using ~1000 samples per dietary condition, and therefore our estimates adequately represent the true population and gene-specific variability.

To compare gene-wise transcript level variability between control and high sugar genes, the MADs and means were ranked within diet and split into 20 quantiles, each containing 438 genes. Then, the overlap between control and high sugar quantiles was calculated by taking the number of shared genes per quantile and dividing it by the total number of genes per quantile (438) and multiplying by 100.

To test for the association between functional features and gene-wise variability in each diet we carried partial Spearman correlation tests between MAD and the continuous features while controlling for potential residual effects of mean expression on expression variability by fitting mean expression level as a covariate. For categorical variables we used Wilcoxon rank-sum tests. The functional and evolutionary features were obtained from a variety of sources. Regulatory DNA features (TATA box, DHS sites, promoter shape broadness, gene and TSS GC content) and functional categories (housekeeping, ubiquitous expression, maternally-transmitted) were obtained from Sigalova *et al.* [14]. Gene lengths and number of alternative transcripts were obtained from FlyBase[15]. Protein-protein interaction connectivity measures (degree, hub, closeness, betweenness, transitivity) were calculated using igraph[16] on StringDB[17] interactions at 0.5 confidence to represent >50% of the total gene set. Transcriptional network connectivity, a combined measure of connectivity from different genomic and transcriptional reporter experiments, was obtained from [18], which in turn, were obtained from [19]. Gene essentiality information was obtained from OGEEv3 [20]. Gene ages were obtained using GenEra [21] and closely-related phylostrata were merged to increase gene number based on author recommendations. dN/dS values were obtained using orthologR on *Drosophila melanogaster vs Drosophila pseudoobscura* [22]. Expression heritability and estimates of average diversity (pi) near the gene body for each gene was obtained from [1].

### Differential expression and variability between diets

To determine whether each gene was differentially variable between conditions, we used a generalised additive model for location, space and shape (GAMLSS)[23], first recommended for differential variability analysis of RNAseq data in [24]. In contrast with other tests for differential variability, like the Levene's and Brown-Forsythe tests, the GAMLSS models the effect of an arbitrary number of variables on the mean and variability of expression (known as the biological coefficient of variation (BCV) the non-Poisson noise or overdispersion) simultaneously so that differential variability can be distinguished from differential mean expression. Further benefits include being able to model untransformed counts under a negative binomial distribution via a log-link function and estimate the explicit effects of covariates on the mean and variability. This test has been shown in two previous publications [25,26] to have a higher true positive rate and lower false positive rate on simulated data compared to other differential variability tests, such as the simpler Levene's test. Following these publications, we show that the results of a Brown-Forsythe[27] on TMM library size-normalised logCPM data with five batches and four SVs regressed out, highly overlap with the GAMLSS results (Fig. S12, table S8).

Three models were defined in GAMLSS; the full model where high-sugar, the five batches and SVs have effects on mean and high-sugar diet has an effect on variability, the full model excluding the high-sugar effect on variance ("null variance"), and the full model excluding the high-sugar effect on mean ("null mean"). The four SVs estimated on the voom() matrix were used rather than the three SVs estimated on the VST() matrix as we needed a representation of the mean effect of the unknown variables on non-variance-stabilised data. It should be noted that SVs and experimental batch effects are modelled to affect only mean transcript level for simplicity and computational time. Statistical significance was tested by a likelihood ratio test of the full model to the null mean model for the mean effect of high-sugar ($\mu$), and the full model to the null variance model for the variance effect of high-sugar ($\sigma$), corrected for multiple testing with the BH procedure. Genes with a FDR <0.05 were called significantly differentially expressed or variable. The models converged after 100 iterations for 8752 (99.8%) of genes at a starting sigma of 0.1, seven genes converged for a starting base σ of 0.5, and five genes failed to converge even if σ was increased until 2. Increases or decreases in mean expression were called by comparing the estimated condition-specific CPM, calculated as $e^{\mu+\ln(1000000)}$. Increases or decreases in variability were called by comparing the estimated condition-specific BCV, calculated as $\sqrt{e}^{\sigma}$.

We downsample the datasets per condition to evaluate first, the effect of sample size in the detection of differentially variable genes, and second to assess whether the slight difference in sample size between control (n = 938) and high sugar (n = 1037) biases the results compared to perfectly even sample size. We find that at least 300 samples per condition are necessary for a stable number of DVGs across replicates for our outbred population, and that the slight difference in sample size across conditions does not bias the results (10).

#### Developmental time and egg-to-adult survival

To follow up on the differential variability results that indicated that development-related pathways had a significantly lower expression variability in the high sugar diet, we set to ascertain the effect that high sugar diet had on two key developmental traits, developmental time and egg-to-adult survival. For this, we used Nex flies from the same outbred population used for RNAseq and whole-genome sequencing, but collected ~150 generations later. Flies from three independent replicate cages maintained on control food were used to seed the experimental vials. These flies were age-matched to be 5-days old and reared in mixed-sex vials until the start of the experiment. 15 vials with 5ml control food, and 15 vials with 5ml high sugar food were seeded with three males and three mated females each. All vials were kept at 25 °C, 65% humidity, and a 12h:12h light:dark cycle. To collect density-controlled egg samples for developmental timing assessments, flies in each vial were allowed to lay eggs for an 18-hour period between 5 PM and 11 AM. Once eggs were collected, two pictures per vial were taken to minimize the effects of light reflections and ensure accurate egg counting. Starting at day four post egg-lay, pupae were counted twice per day until no more pupae emerged.

We modeled the effect of diet on developmental time using a discrete-time survival approach. For each replicate and day, we recorded the number of larvae that completed pupation (count) and the number still not pupated (still_at_risk - count). We then fit a binomial generalized linear model (GLM) with a logit link, specifying the response as the number of pupated larvae out of those still at risk on that day. The model included fixed effects for diet and day (as a categorical variable) to test whether diet affects the daily probability of pupation, accounting for the baseline daily pupation rates. The model can be written as:

$$\mathbf{logit}\,(p_{ij}) = \beta_0 + \beta_1 \cdot \mathbf{Condition}_i + \beta_2 \cdot \mathbf{Day}_j$$

Where condition is either high sugar or control diets, pij is the probability that a larva in condition i pupates on day j, given it has not pupated before day j. Model fitting was performed in R using the glm() function with family binomial. Statistical significance of the diet effect was assessed via Wald tests on the B1 coefficient.

In an independent experiment, egg-to-adult survival was assessed in 12 control replicate and 12 high sugar replicate vials using the same NEX population. Flies were age-matched to be four days old and reared in mixed-sex vials until at the start of the experiment. For each vial, we collected 10 mated females and allowed them to lay eggs for two hours, yielding 80–120 eggs per vial. Then, we discarded the adult flies, then photographed and counted the number of eggs. One replica in control condition was removed from the analysis due to low image quality. Once images were taken, vials were kept at 25 °C, 65% humidity, and a 12h:12h light:dark cycle. Every day from the date of eclosion of the first to the last adult fly (corresponding to day 8 and day 14 after egg lays), adult flies from each vial were collected under $CO_2$ anaesthesia, counted and then discarded. Egg-to-adult survival was calculated for each vial as the ratio of the total number of adults counted to the number of eggs converted to a percentage. The Wilcoxon ranked sum test was used to confirm an effect of the diet on survival. Using Pearson's correlations we showed that the slight differences in egg density in each vial per condition did not bias our results as there is no correlation between the number of eggs and egg-to-adult survival (Fig. S13). Data for both experiments are available in table S12.

#### DNAseq library preparation, data processing, SNP filtering, and annotation

Library preparation, data processing, and SNP calling procedures are described in detail in [1]. In short, raw reads were mapped to the r6.14 *D. melanogaster* genome using BWA[28]. SNP calling was done in GATK. Filters used in GATK's SelectVariant were the following - keep if: ExcessHet<54.69 – corresponds to a probability to violate HW due to heterozygote excess 3.4e-6, biallelic site, present in at least 500 individuals, mean coverage <22x, MQ>50, QD>2.0, FS<60, SOR<3, -2.0<MQRankSum<2.0, -2.0<ReadPosRankSum<2.0). Additionally, given that many of our individual samples have low coverage, we set to NA all individual genotypes with Genotype Quality (GQ) <9 corresponding to a level of confidence of 87.5%. To do this, we first mark these sites using VariantFiltration, and then SelectVariants --set-filtered-gt-to-nocall. Sites with MAF<0.05, present in less than 20% of individuals, and with HW p-adj for heterozygote excess <1e-7 and heterozygote deficit <1e-40 were removed. The resultant SNPs resulted in an LD decay of ~200bp[1].

To obtain a more stringent set of SNPs present across both populations, the following filters were applied within diet, MAF<0.05, genotyped in at least 50% of the samples, and LD<0.8. The filtered SNP sets per diet were then overlapped to retain only common SNPs. This resulted in a set of 383 710 SNPs. This VCF file was finally split by diet and separately reformatted using a combination of vcftools and R commands to satisfy the input requirements for the mapping software. DNAseq quality control metrics of this final SNP set across the population are summarised in Figure S2.

While there is no standard filter on the equality of sample sizes between genotype classes across studies on the genetic basis of variability, a simulation study highlighted that some tests for variability are prone to false positives when (1) the minor homozygote or heterozygote group is small compared to the major homozygote group and (2) the minor allele is variability-increasing [29]. They recommend a minimum heterozygote count of 100. We find that excluding SNPs that fall below this number from the significant veQTL sets (see 'eQTL and veQTL mapping') does not differ from our findings using the full set of SNPs. Specifically, the major trends identified here like the mapping of veQTL for almost all the genes in the genome, their environment-dependent nature (fig. S14), and their MAF bias towards low frequencies (fig. S19), are also recovered with the stringent set of SNPs.

### eQTL and veQTL mapping

To identify the genetic basis of variation in transcript level means and variabilities (Fig. 4A), we carried out both expression quantitative trait loci (eQTL) and variability expression quantitative trait loci (veQTL) mapping.

To identify veQTL, we used squared residual correlation tests implemented in veqtl-mapper [30–32] In these models, the effect of each SNP on mean expression levels is estimated first, and the residuals of that regression are used to estimate the SNP effect on variability by correlation. In this way, genotype-specific variability effects that might be driven by genotype-specific mean effects are accounted for. Alternative ways to account for genotype-specific mean effects that might confound veQTL mapping like *dglm* [33] utilize a joint mean-variance modelling approach instead of a two-step procedure. However, this has previously been shown[34] to be computationally intractable for transcriptome-wide trans-veQTL mapping like our dataset that includes 18 000 genes (9000 genes in each condition) across 400 000 SNPs.

Veqtl-mapper doesn't allow to fit genetic relatedness matrices (GRM) that are critical in outbred populations with related individuals to control for false positives. The only two methods that can model the GRM are *hlmm*[35] and *hglm* [36], where the former is only able to explicitly model 1000 SNPs and the latter is computationally prohibitive in the same way as the *dglm* . We therefore opted for removing the random mean effect of the GRM from the gene count matrix before mapping using GenAbel [37]as previously done in various studies [32,34,38,39]. As the approach relies on statistical frameworks for normally-distributed expression, we began with the logCPM expression matrix with batch and four SVs regressed out. The mean-variability correlation is corrected by procedures highlighted in the following paragraphs. We then split it into diet-specific matrices and did the following for each matrix. First, a GRM per diet was estimated using ibs() on the diet-specific genotype matrix and modelled as a random effect on each gene's expression using polygenic(). Second, for each gene, residual expression (from the grresidual Y field) was obtained from the resulting R dataframe. Then, we applied quantile-normalization using a custom R function as carried out in [32,35,36] to minimize the number of false positives generated by outliers. (see 'defining genotype-by-environment interactions' for a further justification of this choice). Finally, the expression of all genes and all individuals was concatenated to reconstitute a normalised expression matrix per diet. We will refer to this as the GRM-free expression matrix.

It is important to note that there are no methods available to fit genotype-by-environment effects for variability at scale. Accordingly, we run veqtl-mapper for each diet using the GRM-free expression matrices. To map cis-veQTL, we modified the default parameter that detects cis-SNPs in a 1 Mbp window from the TSS, to accommodate the standard 10 Kbp window commonly used to define cis-regulation in *Drosophila* [1,42]. Trans-veQTL were mapped using default veqtl-mapper but with genotype positions recoded to 1:383 710 (the total number of SNPs) to allow every trans-veQTL to be tested. For each gene, we specified the top lowest p-value eQTLs as covariates, up to a maximum of 50 for computational tractability. This approach has been previously shown to account for residual linkage between the focal SNP being tested and SNPs that significantly affect mean expression; such linkage could result in spurious associations (known as 'haplotype effects') between the focal SNP and expression variability[40,43]. As additional stringency would benefit this procedure, we used the top cis-eQTL as defined by the inbuilt 'cis-permutation' mode as opposed to the BH procedure.

We used the Benjamini Hochberg procedure to call significant cis-veQTL and trans-veQTL at a 5% FDR threshold. This was carried out for cis-veQTL by recording the p-values of all cis-tests and using p.adjust() on this complete list of p-values. For trans-veQTL, given the large number of tests, only p-values < 0.0005 were recorded and loaded into R, and following the Matrix eQTL approach [1,34,44], we used p.adjust to apply a Benjamini-Hochberg correction with the partial p-value list and the known number of total tests - *p.adjust*(n=3 361 720 314)

A critical part of our study is the comparison of the molecular and evolutionary properties between eQTL and veQTL. For this reason, we chose tensorqtl[45] for eQTL mapping, which computational speed, computational implementation, and statistical testing framework is the most similar to veqtl-mapper. Tensorqtl, as veqtl-mapper, does not allow to fit a GRM nor GxE interactions in the same model, and therefore both mapping approaches use the same GRM-free matrices, and are conducted within-diet. This allows for a comparison of eQTL and veQTL results that were obtained on the same statistical grounds. The significance of eQTL associations was determined using the same BH approach described above for veQTL.

#### Defining genotype-by-environment interactions

To assess the extent of GxE regulation for genes with eQTL and veQTL as well as the diet-dependent effects of eQTL and veQTL, we used a reciprocal-condition p-value cutoff approach over a range of p-values. We chose this approach given that currently there are no approaches that allow to model GxE interactions for veQTL mapping, as described in the section above. Previously, when other technical reasons prevented us from directly modelling GxE, we showed with empirical and extensive simulations, that this approach behaves well to identify GxE effects of large differential effect between diets, that is, when one QTL has larger effects in one condition and un-detectable effect in the other [46]. It is important to note that the fact that a genetic effect is detected in one diet and not in the other doesn't necessarily mean that the effect is truly zero. It can instead mean that the effect is small enough to go undetected with the statistical approach and sample size used here. We therefore interpret the amount of genetic effects shared by the two diets as a potential underestimation, and discuss the results in that context. However, we note that the sample size used here is well balanced across conditions (938 control and 1037 high sugar), which provides comparable power to detect associations. And, elsewhere we have shown that the number of *Drosophila* genes for which eQTL that can be discovered starts to reach saturation with a sample size of ~1000 individuals [1]. Namely, we defined genes or eQTL/veQTL as 'shared' if the gene-SNP pair had an FDR < 0.05 in the given condition and a FDR < 0.1 in the other condition. A gene/eQTL/veQTL was defined as 'diet-specific' if the gene-SNP pair had an FDR < 0.05 in the given condition and a FDR > 0.1 in the other condition. Due to the large number of tests involved in expression QTL mapping, we evaluated the effect that multiple testing stringency had on the inference of 'shared' and 'diet-specific' effects by fixing FDR 0.05 in one condition, and lowering the FDR in the second condition from 0.05 to 0.2 (Fig. S15; table S14). The general pattern remains the same regardless of which statistical threshold is used for the second condition, with the sole exception of genes with shared-effect veQTL at FDR<0.2. Using a FDR<0.01 minimal threshold rather than 0.05 reduced the number of genes and eQTL/veQTL but likewise did not alter the pattern.

To assess whether the quantile normalisation of the data independently for each condition could drive the GxE results, i.e., introduce differences between CTRL and HS, we repeated the cis-eQTL and cis-veQTL mapping using data that was not quantile-normalised. Given the computational demands, we run this comparison for cis-QTL, but, as the same significance-calling procedure and data normalization was carried out for trans-veQTL, we expect that trans-veQTL will show the same pattern. Across the three comparisons: significant gene-SNP pairs, genes with at least one eQTL/veQTL, and eQTL/veQTL, we find only minor differences and an overall 95% overlap in the results with and without quantile normalization (FDR 0.05 in both conditions) (table S15). This result indicates that the strong environment-dependent results found here are not the result of technical differences introduced by quantile normalization (table S14 and S15).

**Enrichment of eQTL and veQTL in functional SNP features**

To determine whether eQTL and veQTL exhibit different biases in biochemical function we first annotated each of our SNPs with known functional properties. SnpEff[47] was used to obtain the following features – exon mutation, lost start codon, gained start codon, missense mutation, synonymous mutation, 3' UTR, 5' UTR, intron mutation, and intergenic mutation. SNPs were classified as falling into promoters if they were either +/- 100bp within transcription start site locations obtained from the Eukaryotic promoter database [48]. We defined SNPs as being within cis-regulatory modules (enhancers, silencers, others) specific to female adult tissues and transcription factor binding sites based on Redfly location annotations [49]. The significance of the bias relative to non-eQTL and non-veQTL was assessed using a chi-squared test corrected for multiple testing using the Benjamini Hochberg procedure for each combination of control and high sugar, cis- and trans-, eQTL and veQTL.

#### Allele age and direction of variability change

To test if evolutionary recent alleles (derived alleles) of the sets of eQTL or veQTL are biased towards increasing or decreasing effects, we first polarized a greater set of 482 742 SNPs (to include all SNPs used for mapping) following [46] . In short, we used *blastn* in BLAST version 2.10.0 to determine whether a 100-bp region centred on focal SNPs in *Drosophila melanogaster* contained a single orthologous region in *D. simulans* w501(Prin_Dsim_3.0, Genbank GCA_016746395.1) and *D. yakuba* Tai18E2 (Prin_Dyak_Tai18E2_2.0, Genbank GCA_016746365.1) reference genomes. We then aligned unique orthologous regions using MUSCLE version 3.8.31 and called ancestral and derived alleles based on maximum parsimony. We were able to confidently determine ancestral/derived state for 121 683 SNPs out of the 383 710 total used to map eQTL and veQTL.

Next, we calculated for each eQTL-gene and veQTL-gene pair whether the derived allele increased or decreased mean expression or expression variability respectively. To do this, we first identified which allele determines the sign of the estimated phenotypic effect on  a gene's expression outputted by tensorQTL or veqtl-mapper, where a negative sign represents a decreasing effect while a positive sign represents an increasing effect on mean or variability. For eQTL/veQTL where this allele is the derived allele, the assigned effect is retained. For eQTL/veQTL where this allele is the ancestral, the assigned effect is reversed. Because some eQTL/veQTL are pleiotropic (i.e., are associated with more than one gene), we next determined whether the derived allele effect was consistent across all focal genes (i.e., the derived allele always decreases or always increases variability).

For each, trans-eQTL and trans-veQTL sets, within each dietary condition, we estimated the fraction of times the derived allele decreases expression mean or variability, and compared it to that of a null distribution inferred from non-eQTL or non-veQTL (i.e., SNPs with FDR >0.05 for all genes tested). To account for disparities in the number of significant and non-significant QTL sets, as well as for the fact that the effect of eQTL alleles is not consistent across all associated genes, we adopted a downsampling approach where we subsampled 5000 SNP-gene pairs (5 MAF bins with 1000 SNPs each) 1000 times to create derived effect distributions for eQTL, veQTL, as well as null distributions. For each round of subsampling, first, 5000 randomly-selected eQTL/veQTL were selected. Second, a set of 5000 non-eQTL/veQTL SNPs were selected so that their MAF quintile distribution matched that of the eQTL/veQTL set. For each subsample, the fraction of times the derived allele decreased mean expression or variability was recorded. These values were averaged across the 1000 eQTL and veQTL downsamplings to obtain the true value. The 1000 values for non-eQTL/veQTL were used to build the null distributions. ecdf() in R was used to determine which percentile the true values belong to within the null distributions (Fig. 4, E and F; Fig. S20, A and B; table S18). Beyond being computationally simpler than selecting multiple genes per SNP, this approach captures both mean-increasing and mean-decreasing effects of the same SNP across different samples while still allowing us to compute a number that represents the overall bias in the effect of the derived allele.

We further compared the derived allele frequency (DAF) spectrum for either derived alleles with an increasing effect, decreasing effect, or no effect on mean or variability. If the derived allele is also the minor allele, the DAF is the same as the minor allele frequency (MAF). If not, then the DAF is one minus the MAF. We first tested whether alleles with a decreasing effect had a higher median allele frequency than those with an increasing effect using Wilcoxon rank-sum tests (table S19). Then, to test if the DAF spectra for increasing and decreasing eQTL/veQTL differed significantly from that of equivalent-sized sets of non-QTL, we used a slightly different downsampling procedure from before. For each category (control or high sugar, eQTL or veQTL, derived increase or derived decrease), we sampled 5000 MAF-matched QTL and non-QTL 1000 times. For each of the 1000 downsampled distributions, we calculated the median DAF. We also calculated the skewness, which captures how asymmetric the DAF distribution is, where a negative skew indicates a depletion of alleles below 0.5 DAF while a positive skew indicates a depletion of alleles above 0.5 DAF. It captures the switchpoint-like shifts along DAF=0.05 in the full veQTL distributions (Fig. S21, C and D) not seen in the eQTL distributions (Fig. S21, A and B). Wilcoxon rank-sum tests were used to assess if the mean median DAF and skewness differed significantly between the downsampled QTL and non-QTL distributions (table S20).

#### GO term enrichment with g:Profiler

Gene ontology enrichment analyses were conducted using the g:Profiler R package[50]. Significance was determined by hypergeometric tests at p-val < 0.05 following the recommended g:SCS multiple-testing correction. The background gene set corresponded to 8763 genes expressed in *Drosophila* heads for all analyses except the differentially variability analysis, for which the 8752 genes that converged were used.

### Supplementary Figures

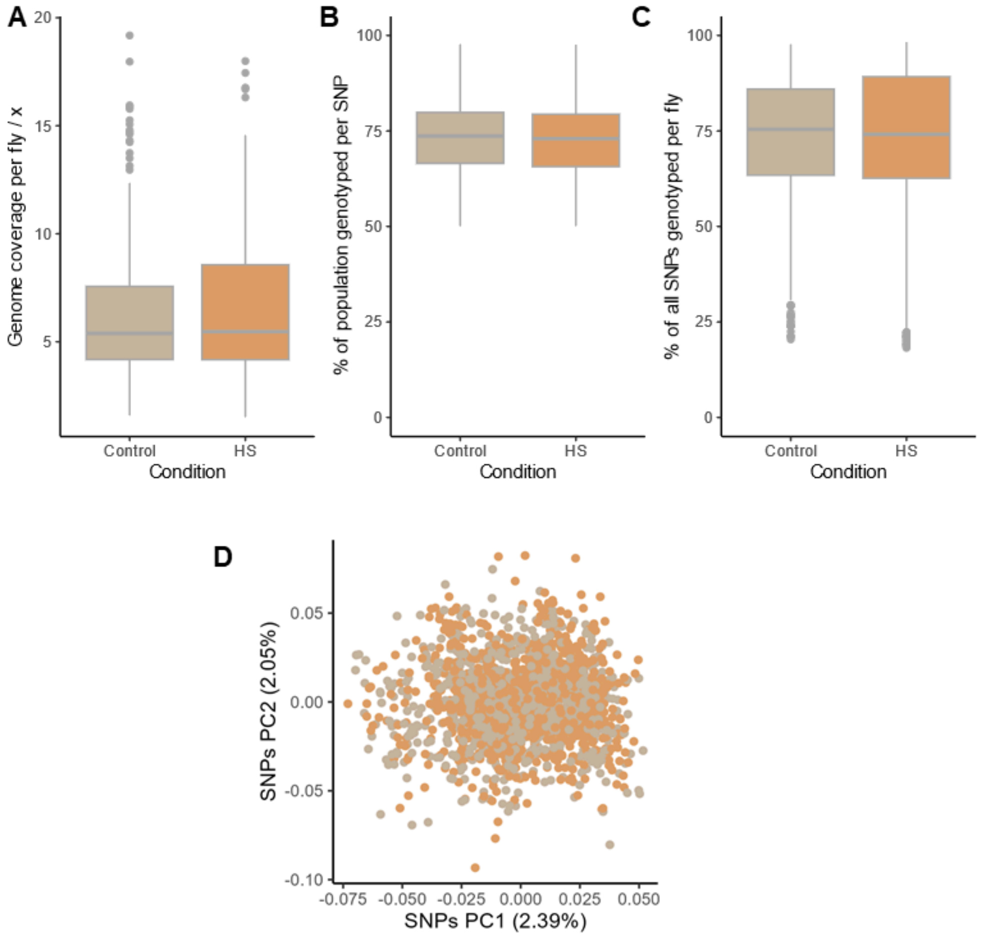


**Figure S1: Properties of RNAseq datasets from two outbred *Drosophila* populations.** (A) Number of reads per sample before mapping and filtering. (B) Transcript counts per fly per condition after after mapping and filtering. (C) Average expression of any given gene per fly per condition. (D) Average expression of each gene in each population. (E) First two principal components of the expression matrix. (C – E) show voom expression, count data subjected to a logCPM transformation and corrected for batch effects and 4 surrogate variables. The Control data (n=938) was obtained from Pallares *et al. Biorxiv*. High sugar (HS) data (n=1037) is presented for the first time here.

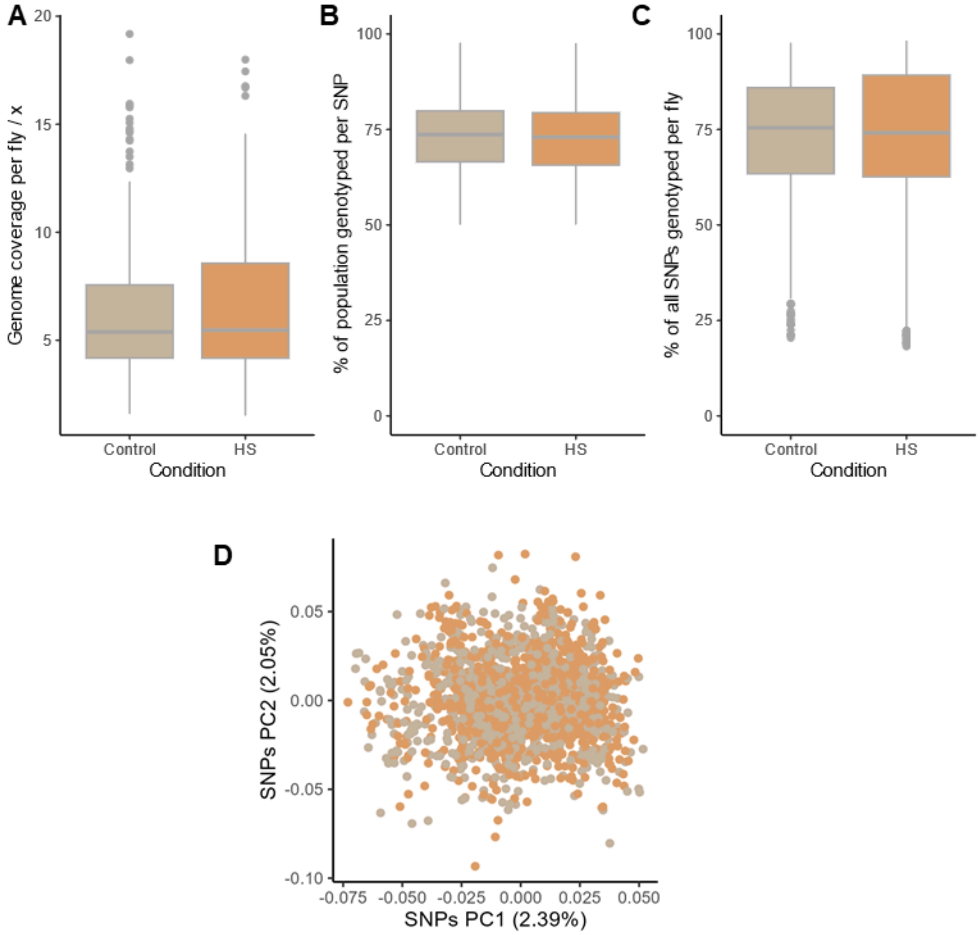


**Figure S2: Properties of DNAseq datasets from two outbred *Drosophila* populations.** (A) Depth of genome coverage per fly per condition. (B) Genotyping rate of each SNP per condition (C) Proportion the SNP set genotyped per fly per condition. (D) First two principal components of the genotype matrix. (B – D) Are for the set of high-quality SNPs used in the analysis (MAF > 0.05, LD R2 < 0.8, genotyped in >50% population). Control data (n=938) was obtained from Pallares *et al. Cell Genomics 2025*. High sugar (HS) data (n=1037) is presented for the first time here.

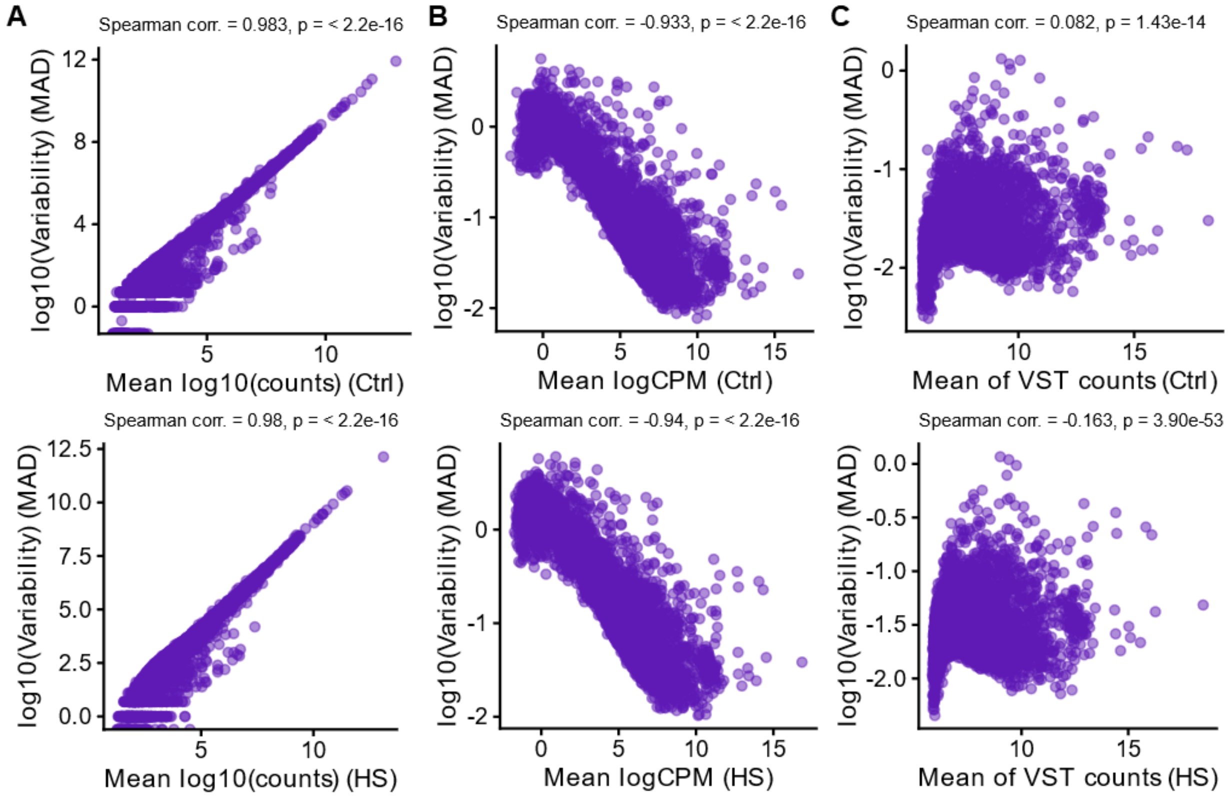


**Figure S3: Mean-variability correlations across different RNAseq data transformations.** The metric of variability used is the median absolute deviation (MAD). Ctrl correlations top and HS correlations below. (A) Per-gene variabilities against means of untransformed raw RNAseq counts yields a strong positive correlation. This is because biologically-replicated raw counts for genes follow a negative binomial distribution with mean and variance ($\mu+\alpha^2$). Both axes are in log scale for easier visualisation. (B) Variabilities against means of normalised RNAseq read counts (logCPM) yields a strong negative correlation. (C) Gene-wise (adjusted) variabilities against means of normalised counts reduces the correlation. MAD of variance-stabilizing transformation (VST) counts shown. Gene-wise variabilities are obtained by first conducting a regression of variability against mean for the whole transcriptome as in (B), then either adjusting the raw counts before variability calculation (*e.g.,* VST MAD) or transforming the variability metric after its initial calculation based on the parameters of this regression (fig. S4). If not applied, all correlations with genomic features will be the same direction as (raw counts) or opposite direction to (logCPM) correlations with means.

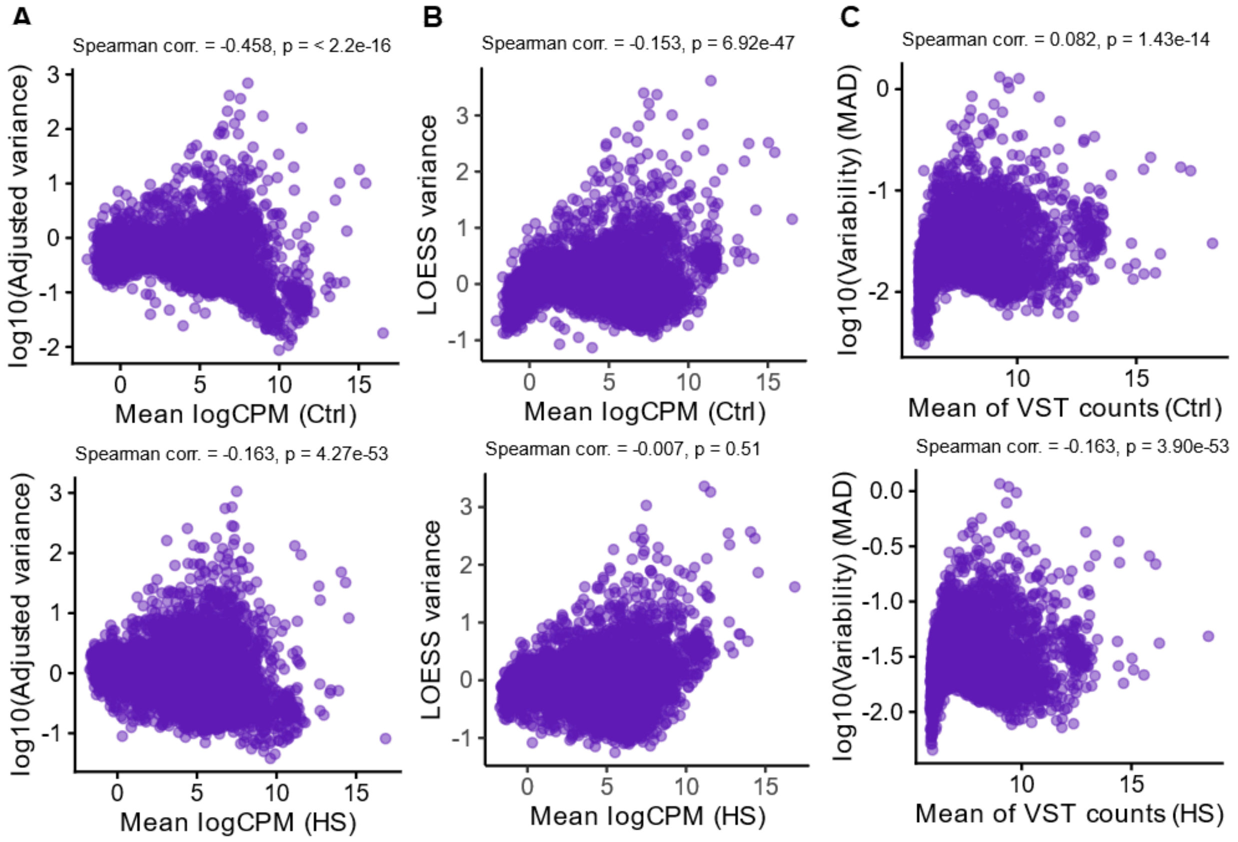


**Figure S4: Comparisons of different adjusted variabilities and their residual correlations with mean in both diets. (**A) Variance of logCPM adjusted by a degree 5 polynomial regression ('adjusted variance') or (B) LOESS regression on the same starting data plotted against mean logCPM. (C) Median absolute deviation against mean counts of VST counts, the variability metrics used in the paper.

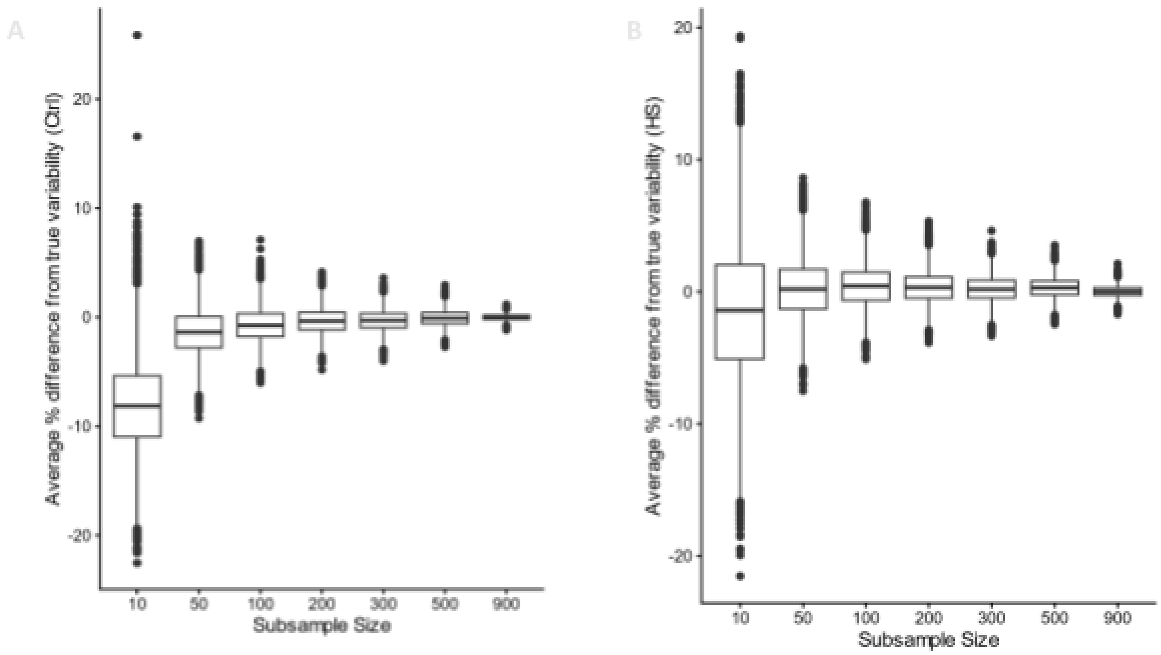


**Figure S5: Downsampling at different sample sizes suggests that the true transcript level variabilities of most genes require 100 samples or more to be obtained under both the (A) control or (B) high sugar diet.** Variability is the MAD obtained from subsamples of VST-transformed counts. Each dot represents the average variability of one gene across 100 samples without replacement. True variability is the variability estimated from the full sample in each condition (n= 938 Ctrl and n = 1037 HS).

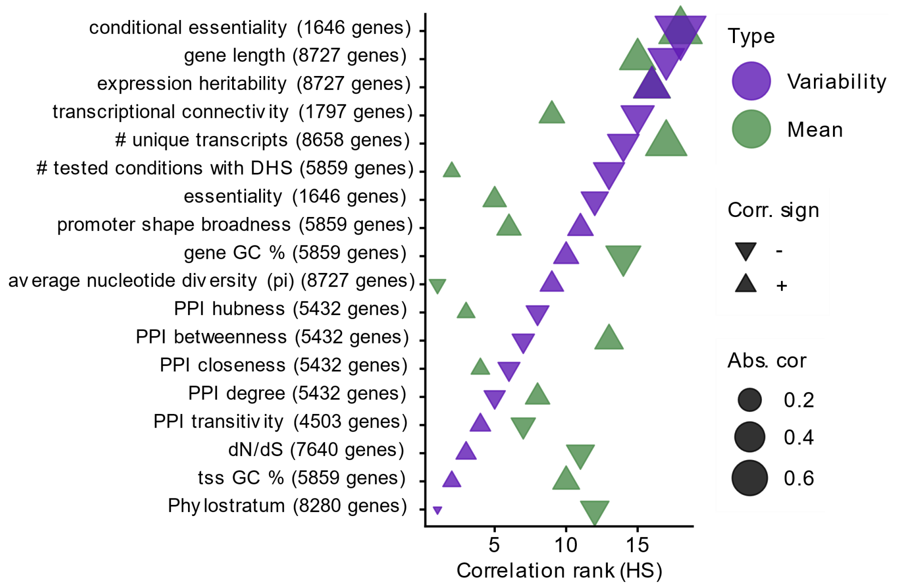


**Figure S6:** Correlates of variability under high sugar. Features are ordered by increasing magnitude and type (red = transcriptional control, yellow = protein function, blue = higher-order function, green = sequence diversity and evolution). Correlations reported are partial Spearman correlations with mean for variability and Spearman correlations for mean. All correlations except Phylostratum for variability are significant (Table S6). Variability is the median absolute deviation of VST counts.

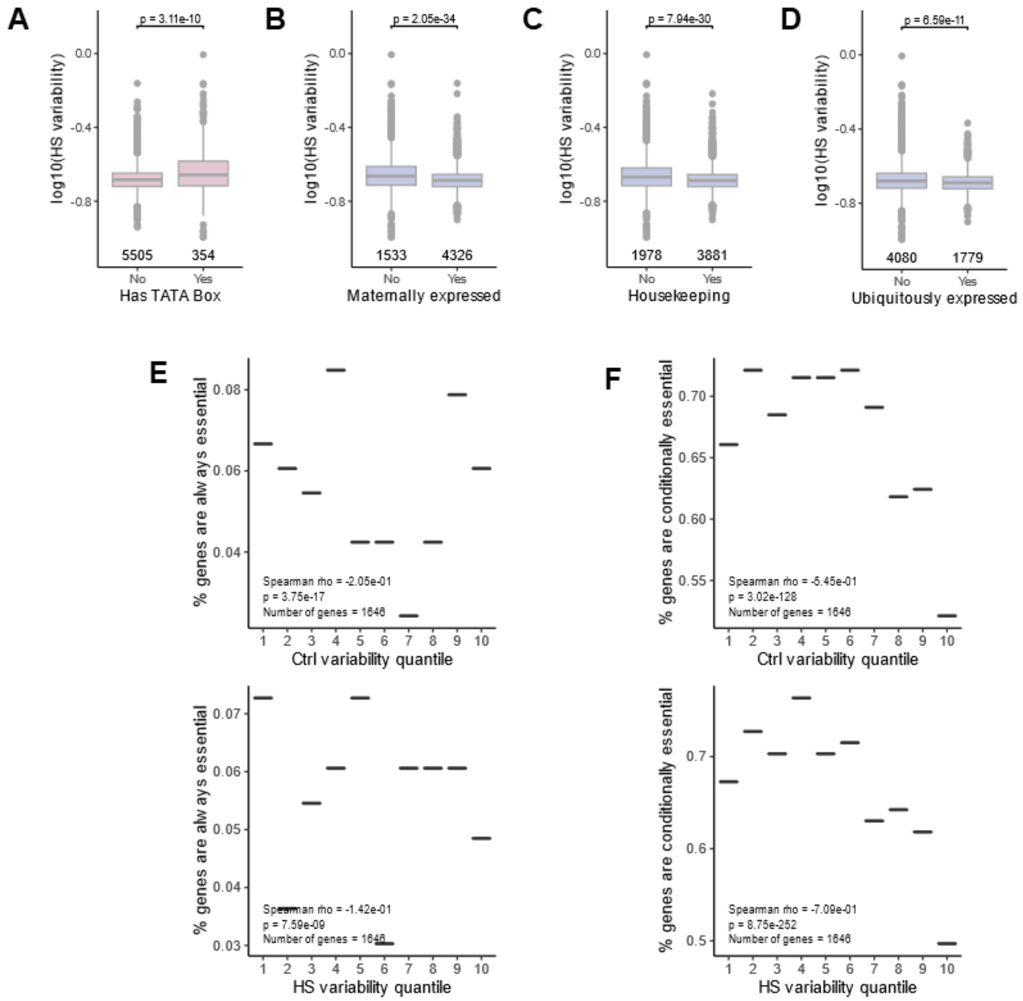


**Figure S7: Categorical functional predictors of gene-wise transcript-level variability.** (A) Genes with TATA boxes have higher variability than those without. Genes that are (B) maternal, (C) housekeeping, (D) ubiquitously-expressed or (E) completely or (F) conditionally essential have lower variability. For (A) – (D), statistical significance was tested using the Wilcox test and boxes are coloured by the category of feature. Partial Spearman correlations were used against redefined variability quantiles for (E-F). Variability is the median absolute deviation of VST counts.

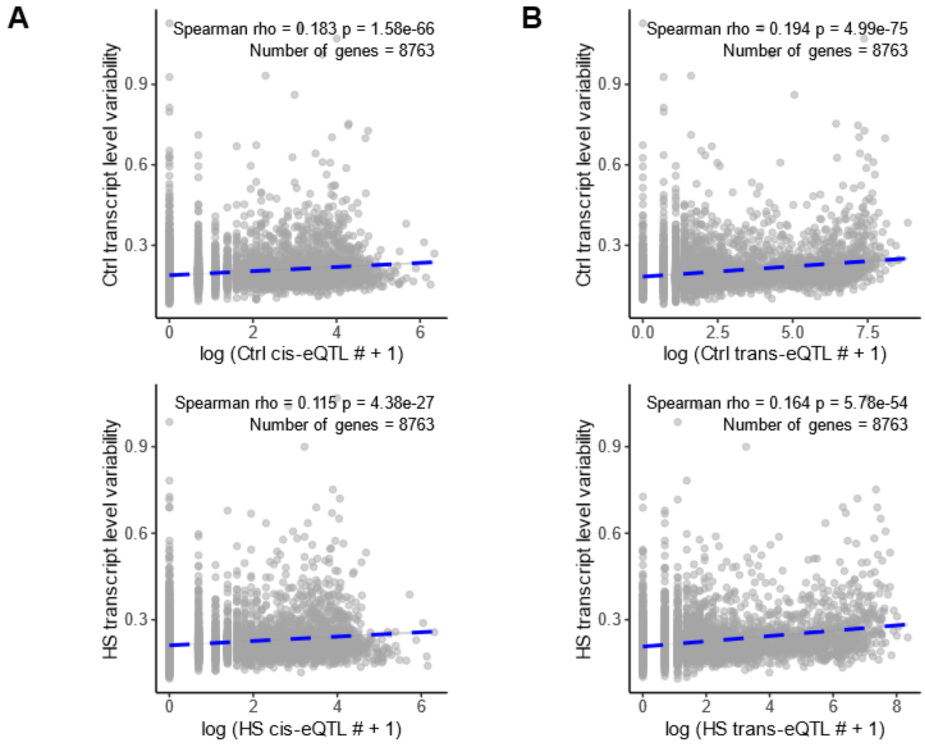


**Figure S8: Number of (A) cis-eQTL and (B) trans-eQTL against transcript level variability.** Blue dashed line is the best fit regression.

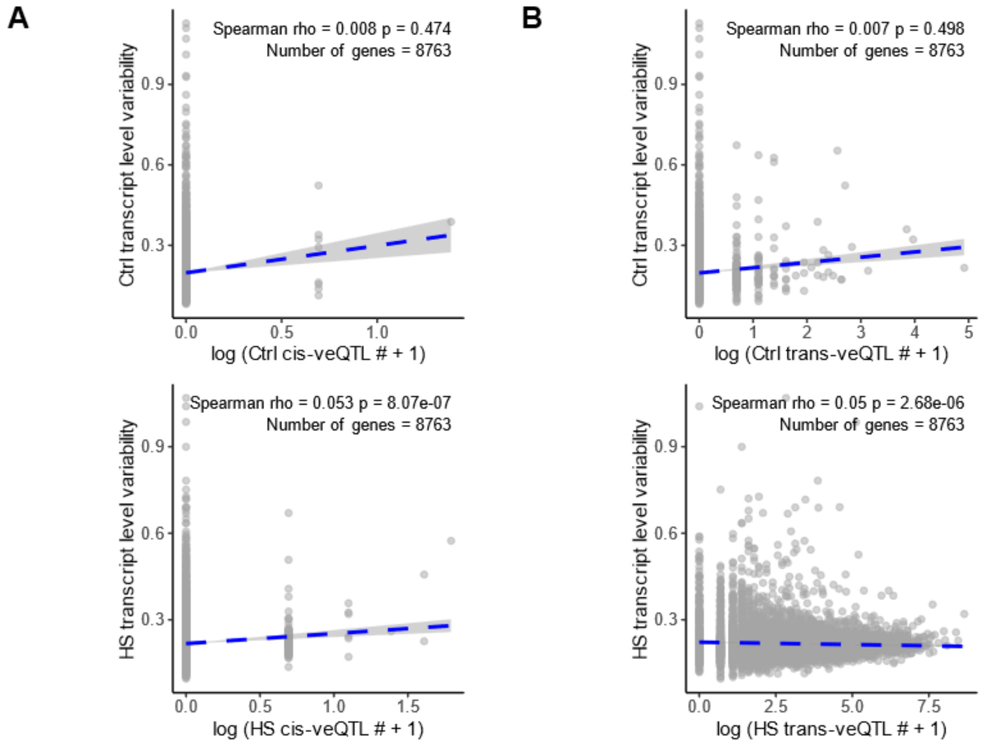


**Figure S9: Number of (A) cis-veQTL and (B) trans-veQTL against transcript level variability.** Blue dashed line is the best fit regression.

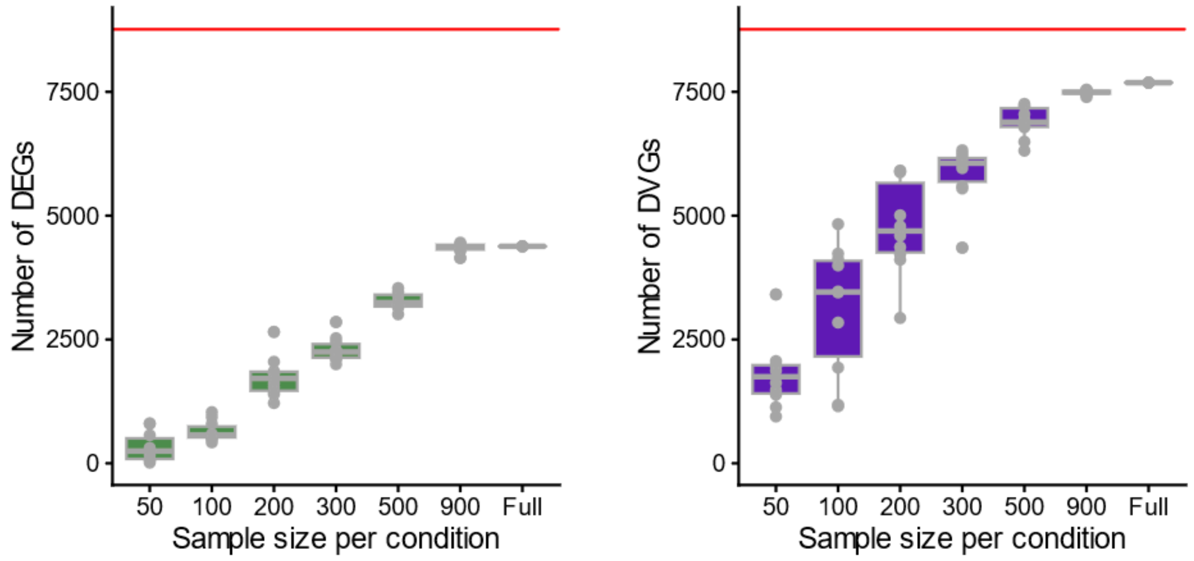


**Figure S10: The effect of downsampling on the number of genes called DE or DV between control and high sugar based on the GAMLSS.** 10 downsamples without replacement performed per sample size. The red line represents the full transcriptome - 8763 genes. The full sample in each condition is n= 938 for control and n = 1037 for high sugar.

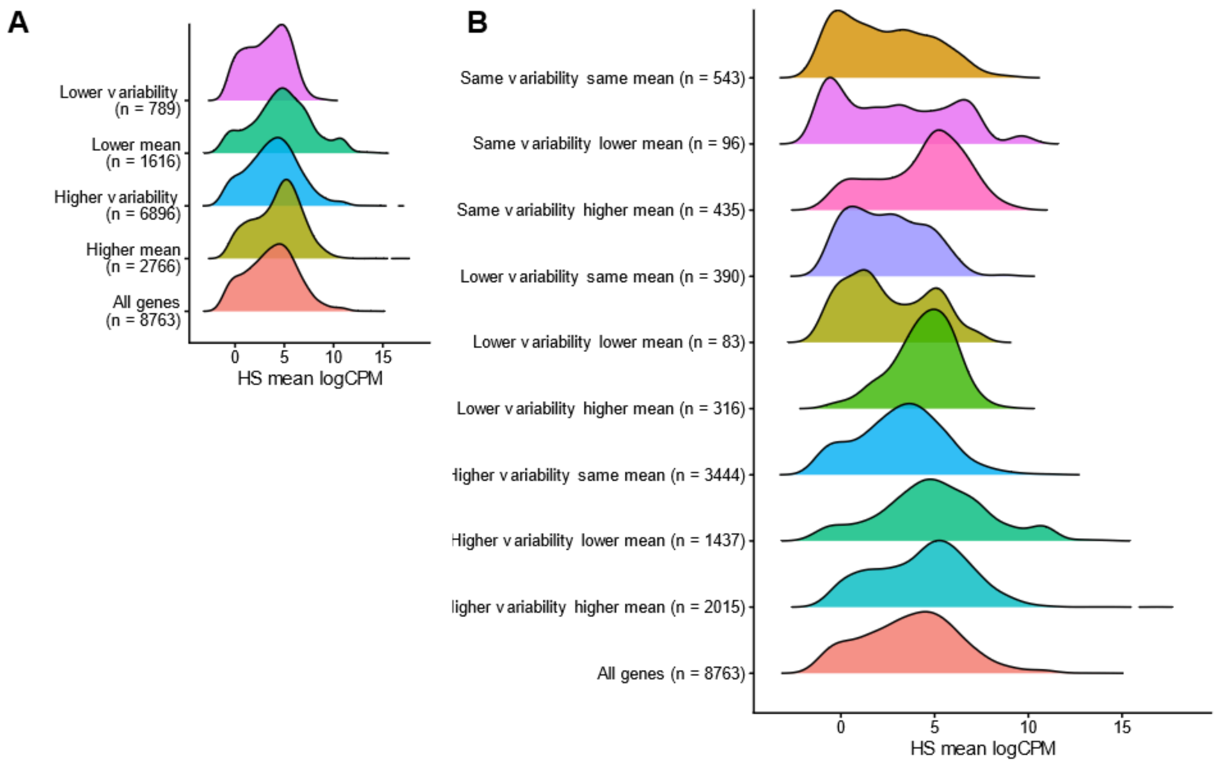


**Figure S11: Distributions of mean logCPM under high sugar of the whole transcriptome.**

(A) DE categories and DV categories. (B) All possible categories of change.

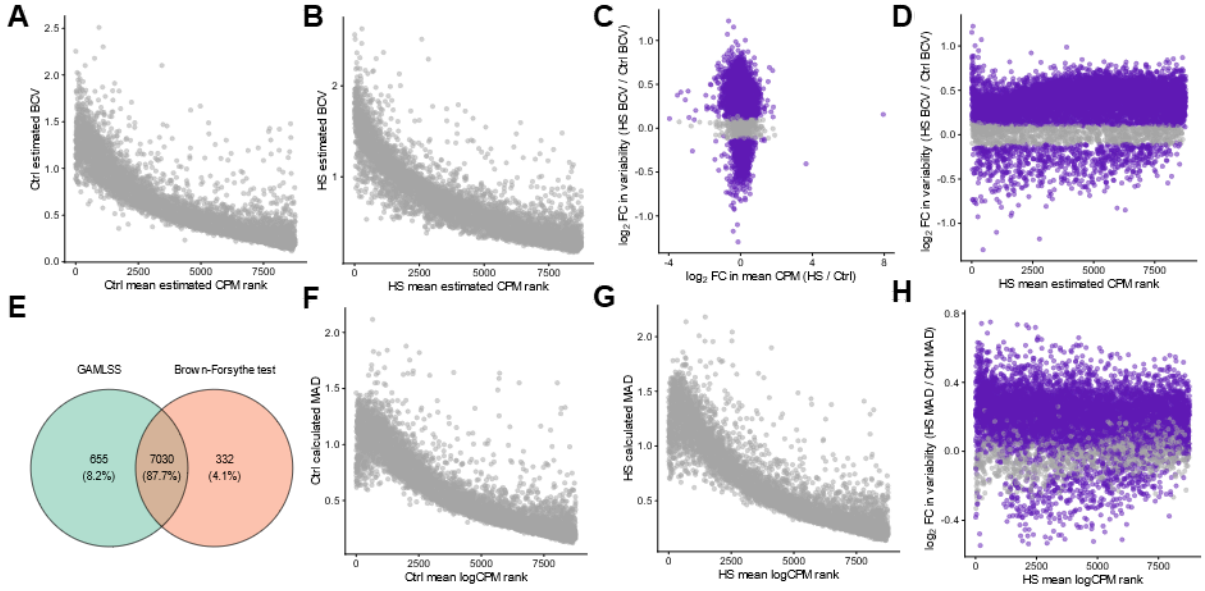


**Figure S12: GAMLSS diagnostic variability-mean plots and comparison with Brown-Forsythe test results** (A) Correlation between Ctrl biological coefficient of variation and mean rank based on GAMLSS model parameters (rho=0.93, p<2.2e-16). (B) Correlation between HS biological coefficient of variation and mean rank based on GAMLSS model parameters (rho=0.93, p<2.2e-16). (C) Correlation between fold change in biological coefficient of variation and fold change in mean between diets (rho=-0.24, p<2.2e-16). (D) Correlation between fold change in biological coefficient of variation and estimated mean (rho=0.17, p<2.2e-16). GAMLSS DVGs are highlighted in purple. (E) Overlap between genes detected as DVGs between GAMLSS and BF test on voom-normalized logCPM data at a BH-corrected p-value of 5%. GAMLSS DVGs are highlighted in purple. (F) Correlation between calculated HS median absolute deviation (MAD) and rank of calculated means based on logCPM data (rho=0.93, p<2.2e-16). (G) Correlation between calculated Ctrl MAD and rank of calculated means based on logCPM data (rho=0.91, p<2.2e-16). (H) Correlation between fold change in calculated MAD and estimated mean (rho=-0.14, p<2.2e-16). BF test DVGs highlighted in purple.

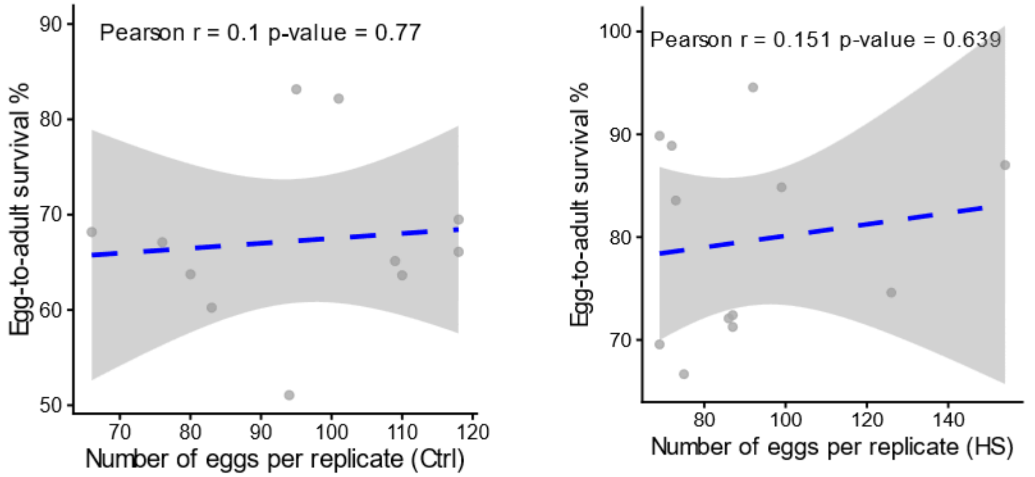


**Figure S13: No significant Pearson correlation between number of eggs and egg-to-adult survival.** Blue dashed line is the best fit regression.

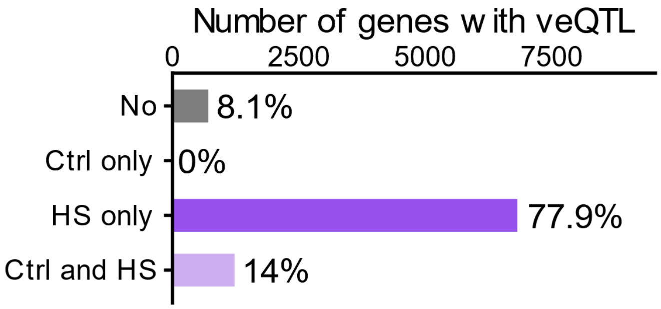


**Figure S14: Context-specificity of genes with veQTL remains after excluding SNPs with heterozygote count below 100.**

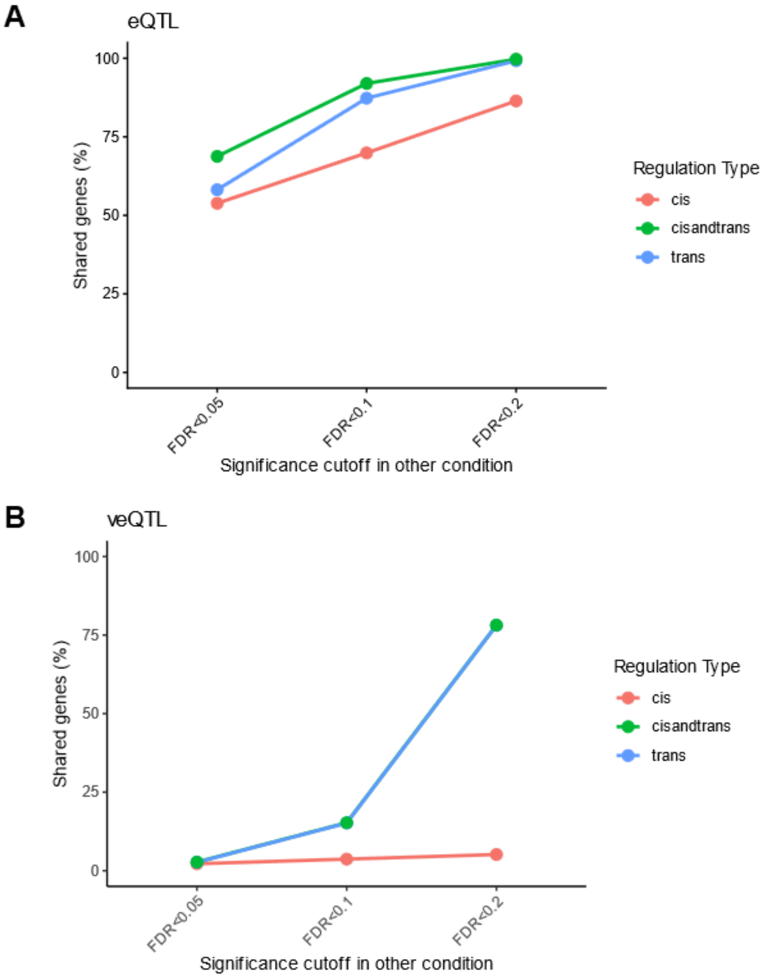


**Figure S15: Genes tend to have condition-shared eQTL but condition-specific veQTL.** The percent of all expressed genes with (A) eQTL or (B) veQTL called as shared between conditions across three cutoff stringencies in the opposing condition.

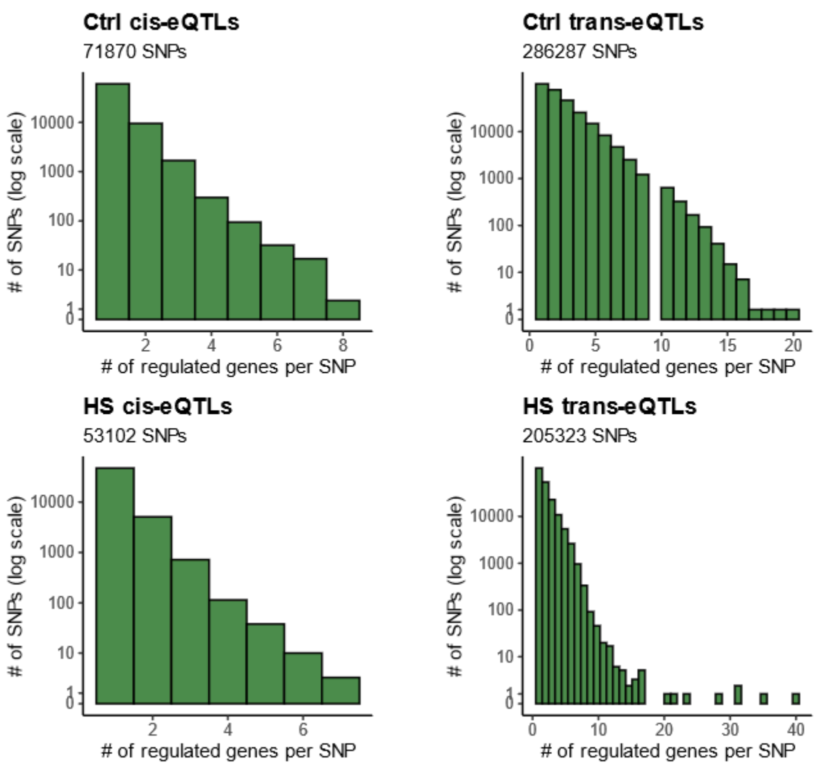


**Figure S16: SNP pleiotropies for eQTL called at FDR < 5% in each condition**

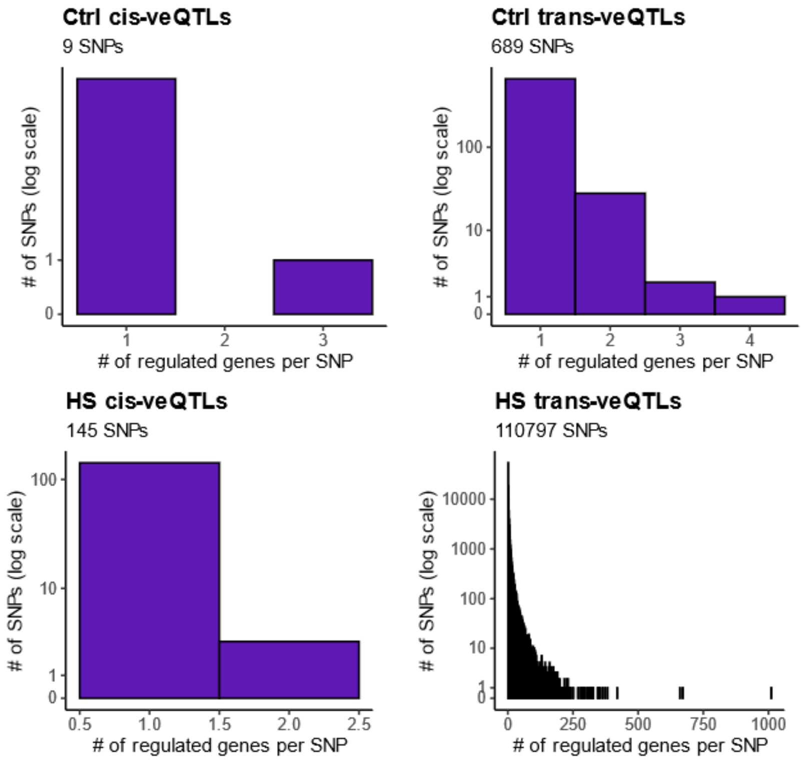


**Figure S17: SNP pleiotropies for veQTL called at FDR < 5% in each condition**

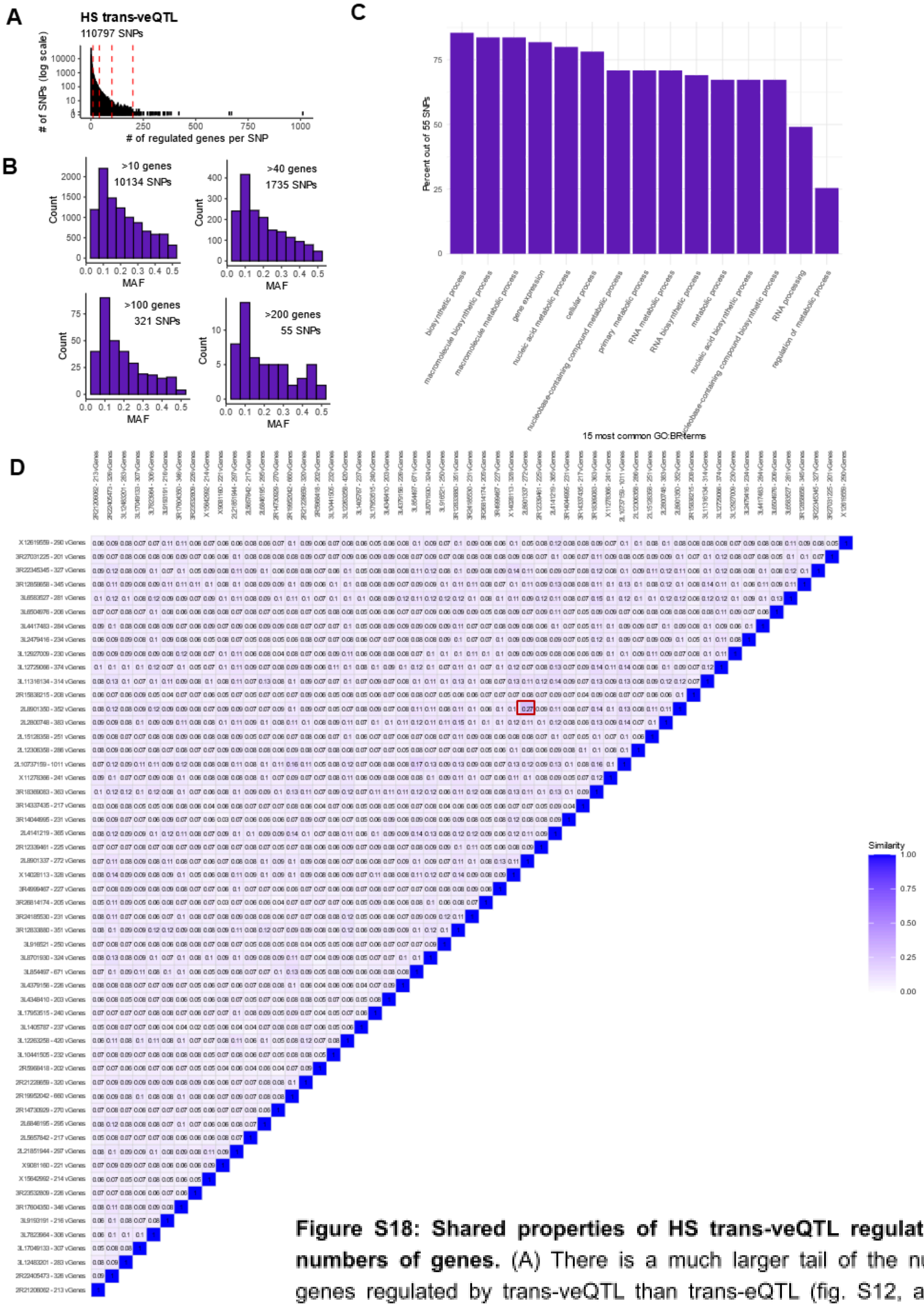


**Figure S18: Shared properties of HS trans-veQTL regulating high numbers of genes.** (A) There is a much larger tail of the number of genes regulated by trans-veQTL than trans-eQTL (fig. S12, and S13). Hypothesizing that these may be exclusively the lowest-MAF veQTL, we first plotted the distribution of MAF for four subsets of veQTL, corresponding to all veQTL regulating >10, 40, 100, and 200 genes respectively (red dashed lines). (B) MAF distributions of veQTL regulating high numbers of genes are biased towards but are not exclusively low MAF. (C) Running GO term enrichment analyses for every gene set regulated by each of 55 SNPs with >200 genes, we find that the most recurrent BPs across the 55 gene sets are related to nucleic acid, and more specifically RNA, synthesis, metabolism and processing. All enrichments by gene set in table S14. (D) Despite this, the percent of genes shared across each veQTL gene set yielded little overlap (27% maximum similarity in red, all other pairs <17%), suggesting each veQTL that affect many genes are minimally functionally interrelated.

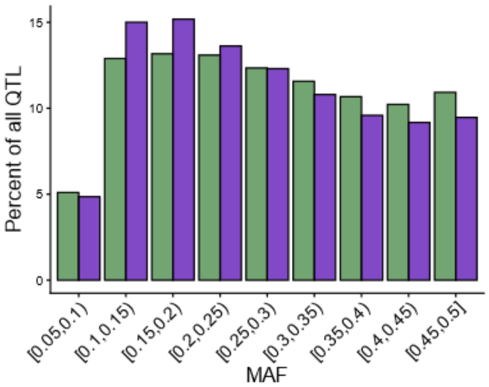


**Figure S19: Lower MAF of veQTL remain after excluding SNPs with heterozygote count below 100.** Minor allele frequency spectrum of veQTL (purple; n=90001, median = 0.256), eQTL (green; n=274105, median = 0.274) shown for comparison (Kolmogorov Smirnoff test p<2.2e-16, Wilcox text p< 2.2e-16).

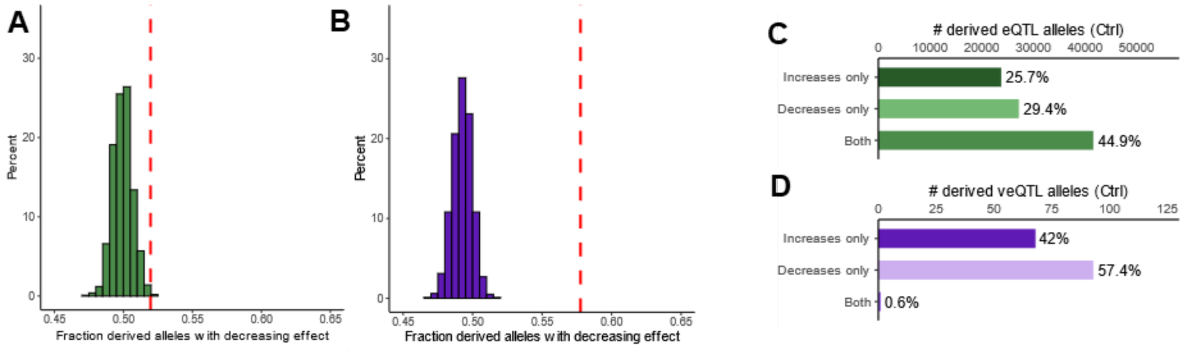


**Figure S20: Effect biases of derived alleles at control trans-eQTL and trans-veQTL.** (A-B) Fraction of subsampled QTL with a decreasing effect (red dashed line represents the median fraction) compared to subsampled non-QTL (histograms). 1000 subsamples of 5000 MAF-matched SNPs each were used to account for large imbalances between the numbers of QTL and non-QTL. For Ctrl veQTL, the fraction of the full SNP set is represented instead due to low number (table S18). (A) 51% of derived alleles of eQTL decrease mean expression, representing the 0.002 percentile of the non-eQTL distribution (green). (B) 57% of derived veQTL alleles of eQTL decrease variability, lying outside the distribution of non-veQTL (purple). (C) Derived alleles at eQTL can only increase, only decrease, or gene-depedently increase or decrease the mean expression of the genes they regulate. (D) Derived alleles at veQTL tend to only increase or decrease the variability of all genes they regulate.

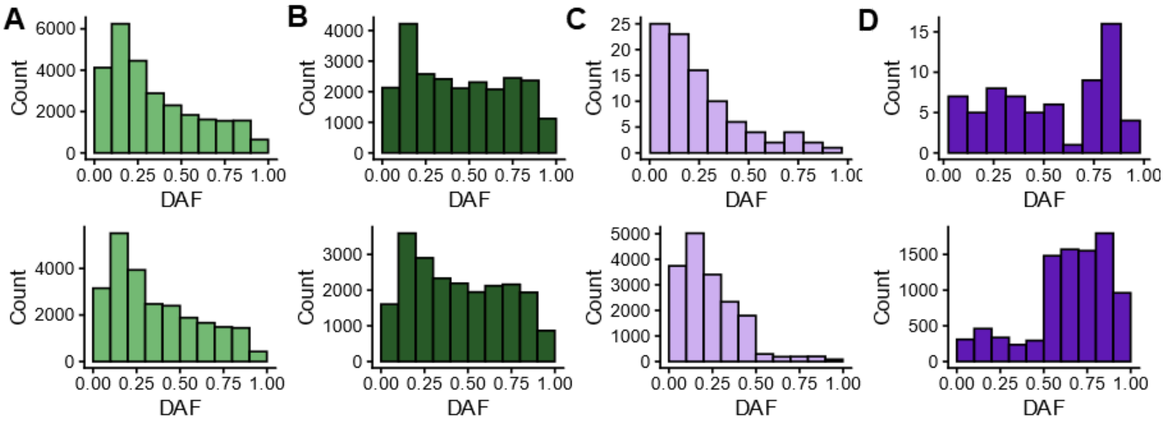


**Figure S21: DAF of alleles at trans-eQTL and trans-veQTL that increase or decrease mean or variability.** Top is control while bottom is HS. (A-B) DAF of eQTL where the derived allele exclusively (A) decreases and (B) increases mean. (C-D) DAF of veQTL where the derived allele exclusively (C) decreases and (D) increases variability only. The median DAFs are significantly different between increasing and decreasing sets. (Wilcoxon ranked sum test p-value < 2.2e-16, Table S20). DAF distributions of downsampled QTL are significantly different in median DAF and skewness than downsampled non-QTL (Wilcoxon ranked sum test p-value < 2.2e-16, table S21).
